# The evolutionary genomics of incipient endosymbiosis in wild rhizobia bacteria

**DOI:** 10.64898/2025.12.03.692210

**Authors:** Angeliqua P. Montoya, Kyson Jensen, Joel S. Griffitts, Stephanie S. Porter

**Affiliations:** School of Biological Sciences, Washington State University, Vancouver, Washington, 98686; Department of Microbiology and Molecular Biology, Brigham Young University, Provo, Utah, 84602

## Abstract

The advent of endosymbiosis underlies evolutionary innovation and ecosystem function. However, whether free-living partners tend to benefit or exploit each other during incipient endosymbiosis remains a dilemma. Rhizobia bacteria are plant endosymbionts capable of initiating root nodules and fixing nitrogen due to genes carried on mobile genetic elements (MGEs) such as the symbiosis island (SI). We conjugated marked SIs into the genomes of nonnodulating strains, which was sufficient to generate *de novo* root nodule-forming endosymbionts. Most novel endosymbionts originated as commensals that incurred no detectable costs to host plants, in contrast to predictions of exploitation. In fact, a third of endosymbionts originated as nitrogen fixing mutualists. Consistent with phylogenetic limits to transfer of MGE function, novel endosymbionts derived from more closely related SI donor and recipient strains showed greater nitrogen fixation. However, we did not detect phylogenetic limits to SI transmission, which could reflect selfish selection for generalized horizontal transfer of this MGE. In fact, the SI was able to displace other genomic elements residing at its characteristic tRNA gene insertion site. We thus provide genetic, genomic, and functional evidence of how MGEs can potentiate and constrain major evolutionary transitions to expand bacterial niches, with cascading effects on host organisms.

## INTRODUCTION

The advent of endosymbiosis underlies evolutionary innovation and ecosystem function across diverse systems (1–3). From N_2_ fixing rhizobia to photosynthetic zooxanthellae, cells that evolve to live within the cells or organs of a host organism are key members of plant and animal microbiomes that have far reaching effects on diversity and ecological interactions (4,5). However, initial evolutionary steps toward endosymbiosis from free living lineages are often unclear. This is especially true with regard to the genetics of early niche expansion into endosymbiosis, because subsequent evolution can obscure initial events in this transition (1,6,7).

While many endosymbioses are mutually beneficial, endosymbiosis may not initially arise as a mutually beneficial interaction. If the evolutionary origin of endosymbiosis were to occur via *de novo* mutualism, this would require that loci that confer initial endosymbiosis benefit both partners and that initial partner fitness alignment is maintained such that mutations that increase one partner’s net fitness are beneficial or neutral to the other (1,8). More simply, endosymbiosis could originate as an exploitative or commensal interaction. Here, loci underlying the origin of endosymbiosis could be selected to increase one partner’s fitness without regard to fitness effects on the other (9–11). Subsequent evolution during which a host preferentially allocates resources to superior partners in the interaction, for example via host-imposed control (12), could later select for mutually beneficial mutations (1,13). Understanding which of these partner fitness conflict and alignment outcomes occurs during incipient endosymbiosis is a critical frontier with broad implications for microbial niche evolution and our ability to engineer synthetic endosymbiosis (14,15,15,16).

Loci that expand microbial niches often reside within mobile genetic elements (MGEs) that can move between genomes via horizontal gene transfer (HGT) (17). MGE transmission can enable locally advantageous loci to spread across distinct bacterial lineages during genespecific sweeps in response to local selection (18). MGEs are also broadly implicated in microbial transitions from free-living lifestyles to endosymbiosis (19–21). Such "symMGEs" are often carried by only a fraction of the lineages in a population possibly because they confer local rather than global fitness benefits and can incur costs due to MGE replication, expression, and transfer mechanisms (22,23). Lineages carrying symMGEs may serve as a reservoir from which a potentially vast population of non-endosymbiotic lineages could acquire the MGE (21,24). If symMGEs are regularly transmitted through populations, microbial lineages have the potential to undergo major lifestyle shifts as they gain and lose them. However, the extent to which genetic interactions (i.e., epistasis) constrain the free transmission and function of symMGEs in microbial populations remains an open question.

The costs of symMGE carriage could select for adaptations to either ameliorate these costs or to prevent symMGE uptake (25,26). On one hand, replicons bearing a symMGE can accumulate compensatory mutations that minimize symMGE costs and/or enhance functional benefits (13,27,28). Such specialization or co-adaptation between symMGE and chromosomal replicons can result in *higher* symMGE transmission and function for donor and recipient genomes that are more closely related (22,29), or ecologically or geographically proximate (30–33). On the other hand, the high cost of symMGEs carriage can also select for defense systems that inhibit foreign MGE entry or maintenance. Here, historical opportunities to evolve defenses against MGEs could lead to *lower* symMGE transmission and function among donor and recipient genomes that are more closely related, or more proximate (34,35). Understanding which of these contrasting scenarios occurs in natural populations is critical to understanding how microbial traits evolve during the origins of endosymbiosis.

To probe transmission and early fitness effects during the transition to endosymbiosis, we quantify sources of genetic variation for transfer and function of a symMGE in rhizobia soil bacteria. In rhizobia, symMGEs bear loci necessary for N_2_ fixing intracellular endosymbiosis within the root nodules of legume plants. Rhizobia symMGEs can be transferred into nonsymbiotic rhizobia, or even non-rhizobia (24,36–38). Acquisition of a symMGE can transform free living bacteria into endosymbionts capable of nodulation and sometimes even N_2_ fixation, though transfer may not succeed and some transfers confer no endosymbiosis function (24,36,37,39,40). We examine *Mesorhizobium* rhizobia that differ in the presence or absence of a symMGE, the symbiosis island (SI) to reveal the evolutionary genetics of transmission. These *Mesorhizobium* were isolated across natural Western USA landscapes from the root nodules of native host plants (41,42) and include serpentine ecotypes that adapt to heavy metal stress via a metal tolerance MGE, as well as non-serpentine ecotypes that typically lack the metal tolerance MGE (41,43).

To investigate the genetics and genomics of partner fitness conflict and alignment during incipient endosymbiosis we quantify sources of evolutionary potential for the transition to endosymbiosis. We tracked SI transmission by inserting a selectable marker into the SI of nodulating donor strains and mating them with SI- strains incapable of forming root nodules. To test for genetic and epistatic constraints on SI transmission, we first asked, 1) Can we induce SI transmission in mating trials and is the propensity for transmission genetically variable? We succeeded in SI transmission across many genotypes and sequenced the resulting transconjugants to ask, 2) What are the genomic outcomes of novel SI acquisition? We then inoculated transconjugants onto host plants to investigate, 3) Are novel endosymbionts mutualists, commensals, or antagonists? SI acquisition most commonly generated nodulating commensal endosymbionts though *de novo* mutualists also emerged. We investigated the genetics of these effects by evaluating, 4) How does donor-SI vs recipient genotype impact partner fitness outcomes in novel endosymbiosis? Upon finding heritable variation in both SI transmission and symbiotic function, we asked, 5) Do phylogenetic, geographic, or ecotypic factors predict SI transmission and symbiotic function?

## METHODS

### Overview

To investigate the genetics, genomics, and functional effects of transmission of the *Mesorhizobium* symbiosis island (SI), we inserted a selectable marker into donor SIs and investigated SI transmission in mating trials. We mated 30 SI+ donor strains and 6 SI- recipient strains in 180 pairwise mating trials and used genome sequencing and colony PCR to confirm SI transfer for a subset of transconjugants. To evaluate the fitness consequences of SI transmission, a subset of transconjugants and their respective SI donors were inoculated onto host plants and fitness and functional traits were measured for both novel endosymbionts and hosts (Figure 1).

**Figure 1.**
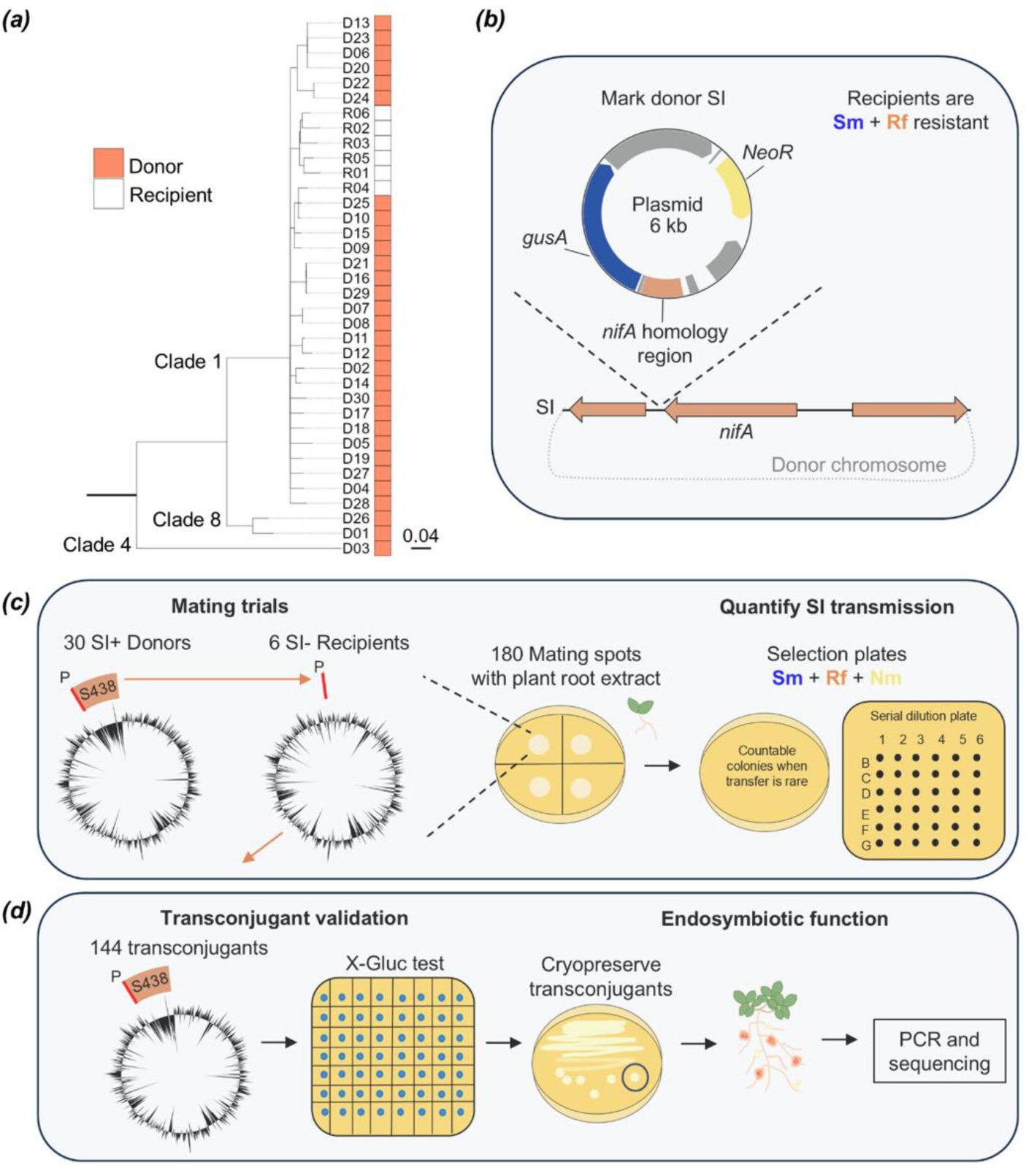
Experimental SI transfer, selection of SI gain transconjugants, and test of symbiotic function. **(a)** Phylogeny of donor (pink) and recipient (white) strains based on 345 conserved amino acid sequences (90) and constructed with RaxML (91). Nodes have support values >70. Scale bar is amino acid substitutions per site. **(b)** A plasmid containing selectable marker genes (beta-glucuronidase (*gusA*); neomycin resistance (*NeoR*)) was integrated into the donor’s SI via homologous recombination (Fig. S1). The recipient strain is resistant to streptomycin (Sm) and rifampicin (Rf). **(c)** 180 pairwise matings occurred within spots of donor and recipient cells. Shown is a marked SI in a donor genome and the Phe-tRNA gene (P; red and enlarged for visualization) SI insertion site in a recipient genome, and genome-wide GC content (black). Two selection plating methods quantified SI gain transconjugants on Sm, Rf, and Nm supplemented antibiotic plates. **(d)** SI gain transconjugants were blue on X-Gluc media due to *GUS* activity and were cryopreserved. To evaluate endosymbiosis, 15 SI gain transconjugants and SI donors were inoculated onto plants. SI transfer and identity were confirmed for a subset of transconjugants via genome sequencing and PCR.

### Symbiosis island donors and recipients

To track transfer of the *Mesorhizobium* symbiosis island from SI+ donor strains to SI- recipient *Mesorhizobium* strains, 30 SI+ donors were transformed with a marker plasmid. The plasmid contains genes for neomycin (Nm) resistance and beta-glucuronidase (*GUS*), which cleaves a substrate so cells appear blue in the presence of X-gluc (5-Bromo-4-chloro-3-indoxylbeta-D-glucuronide). The plasmid also contains a region of homology with the transcriptional activator for the nitrogenase complex, *nifA*, such that the plasmid can recombine into the SI (Supplemental Information; Figure S1). One additional donor strain (not of the 30 donors) was mislabeled, so this strain and products stemming from it were removed. Six recipient strains that lack the SI were originally isolated from root nodules (44), and contain the Phe-tRNA gene required for SI insertion, were selected to be resistant to streptomycin and rifampicin (45). Two recipients (R05 and R06) contain putative genomic islands (GIs) already inserted at the Phe-tRNA with a 17-bp direct repeat of the Phe-tRNA gene at the end of the GI, while four recipients have no predicted GI at this Phe-tRNA gene, and one recipient (R02) has two Phe-tRNA genes. *Mesorhizobium* strains in this study (Table S1) were incubated at 28°C on Tryptone Yeast (TY) agar (46) and as appropriate were supplemented with 100 μg/mL neomycin (Nm), 200 μg/mL streptomycin (Sm), 100 μg/mL rifampicin (Rf), and/or 50 μg/mL X-Gluc.

### Symbiosis island transfer

To transfer the SI between *Mesorhizobium* strains, 30 donor and 6 recipient strains were bi-parentally mated in all pairwise combinations (180 matings). Given the large number of matings, we split the SI transfer matings into two assays (details in Supplemental Information). Donors and recipients were grown on respective antibiotic supplemented TY agar for 4 days, then cells were rinsed and suspended in TY broth and normalized to 6 × 10^8^ cells mL^−1^ using the formula, cells mL^−1^ = OD_600_ × 5.8 × 10^7^ (47). Mating spots consisting of 10 μL donor cells and 10 μL recipient cells, or each strain alone as negative controls, were mixed with 10 μL of plant root extracts and plated on TY agar and incubated for 40 hours. We included aqueous root extracts from native host legumes in the mating spots because these can stimulate transfer of symbiosis gene elements in rhizobia and would occur naturally at likely sites for SI transmission (48,49). We germinated two *Acmispon wrangelianus* and two *A. brachycarpus* genotypes (8 seeds per genotype), macerated roots in 4 mL water, filter sterilized, and stored root extract at 4°C in the dark.

### Quantifying symbiosis island transmission

At the end of incubation, each mating spot was suspended in 350 μL of TY broth with Nm. We quantified the number of transconjugants for each mating and control spot using two selection plating approaches on TY agar supplemented with Nm, Sm, and Rf: 100 μL of the mating culture was spread on 100 mm plates, and 50 μL of the mating culture was serially diluted and plated in droplets on square 120 mm plates. Each plating approach was conducted with three technical replicates that were incubated until single colonies formed and colony forming units (CFUs) were counted. Here, transconjugants showed anticipated antibiotic resistance profiles and their count comprised our measure of SI transmission, *transconjugants per mating*. Cells from each of the 100 mm selection plates with growth were transferred to TY agar supplemented with X-Gluc and Nm. Here, blue cells were considered to be transconjugants and a subset were reisolated as single colonies, cultured, and cryopreserved.

### System confirmatory PCR and sequencing

A subset of transconjugants were tested using colony PCR and sequencing to confirm SI presence and the identity of the recipient genotype. In PCR assays, we used genotype-specific primers to confirm recipient genotype and we used established primers to amplify part of the *nodA* gene in the SI (44) to confirm SI presence in 10% the transconjugants (see Supplementary Information; Table S2).

To confirm: 1) integration of the plasmid marker into the donor SI, 2) SI acquisition in transconjugants, and 3) identity of recipients and respective transconjugants, a subset of the donor and transconjugant strains were genome sequenced using Oxford Nanopore Technology by Plasmidsaurus with custom analysis and annotation. We used 5 donors and 15 transconjugants in a greenhouse common garden experiment (below) and sequenced these strains (Table S3). We also used 5 donor and 2 recipient genomic sequences from Kehlet-Delgado et al. (2024) and 4 recipient genomic sequences from Montoya et al. (2025). Sequence alignments were performed between the wild-type donor and SI-marked donor or respective recipient-transconjugant pairs using *progressiveMauve* (50), pairwise average nucleotide identity (ANI) was calculated using FastANI (51) and genomic maps were visualized in proksee (52).

### Symbiotic function of SI gain transconjugants

To evaluate the consequences of SI transmission for symbiont and host fitness, we grew 5 SI donors and 15 transconjugants in symbiosis with a single genotype of native *A. wrangelianus* (SM2A9) in single-strain inoculations. To compare each donor to a few transconjugants, the transconjugants are derived from 5 SI donor genotypes, each transferred into 3 recipient background genomes. Strains were selected haphazardly except we chose to include two donors that were more phylogenetically distinct from the other strains to achieve phylogenetically generalizable findings. We inoculated these 20 strains and a no-rhizobia control treatment onto host plants across 11 randomized complete spatial blocks for a total of 231 plants in the Washington State University Vancouver greenhouse (45.7328054° N, 122.635967° W).

Seeds were scarified, surface-sterilized with chlorine gas for four hours (44), imbibed in water overnight, and stratified on 1% agar water plates in darkness for 10 days at 4°C. Seeds were planted on 13 October 2023 in 164 mL pots (SC10R ‘Cone-tainers’, Stuewe and Sons Inc., Tangent, Oregon, USA) filled with an autoclaved 1:1 mix by volume of potting soil (Sunshine Mix #1, Sungro Horticulture, Agawam, MA) and inert quartz sand, with a 30 cm cotton wick. Seeds germinated and established for 6 weeks in a greenhouse with 14 hour, 18°C days and 15°C nights, and ultra-fine mist irrigation (44,53).

*Mesorhizobium* strains were grown on TY agar for 4 days, rinsed, and resuspended in water at 1 x 10^7^ cells per 900 μL (based on OD_600_) (44,47). To contain transgenic inocula, we halted mist irrigation and placed each pot in a 175 mL test tube containing 75 mL of sterile water in contact with the wick. Each plant was inoculated with 900 μL of inoculum or a sterile water control. Plants were fertilized with 1 mL of nitrogen-free Fahraues (54) at 2- and 5-weeks post inoculation. Water was replenished as needed. Blocks of plants were harvested starting 29 January 2024 with an average of 2 blocks per day.

We measured multiple response variables in the greenhouse experiment (*in italics*). At harvest, shoots were clipped and roots were washed and stored at 4°C. Plant fitness was measured as dry *shoot mass*. Nitrogen fixation was estimated as percent nitrogen (*%N*) in apical leaf tissue by dry mass from 6 blocks (Stable Isotope Core Laboratory, Washington State University, Pullman, WA). One leaf sample was removed because nitrogen was below detectable levels. We measured multiple proxies for rhizobium fitness: the *number of nodules* on a plant (*pink* and *white*), *nodule width* for the largest nodule per plant, and *CFU/nodule*, the number of progeny within the largest nodule for each plant from 5 blocks. We considered white nodules to fix low or no nitrogen or be immature, and pink nodules containing leghemoglobin to be *N-fixing nodules* (55,56). To calculate colony forming units (CFUs) per nodule, nodules were surface sterilized, crushed, and six serial dilutions were drop plated on TY agar in three replicates (47). Extracted nodules from blocks 1-5 and the remaining plant roots from blocks 6-11 were photographed, and nodule diameter was measured in image J (57). We reisolated 13/15 transconjugants from nodules following Kehlet-Delgado et al. (2024). However, we were not successful for 2 transconjugants (R02^SI_01^ and R03^SI_03^) despite several attempts.

### Statistical analyses

All analyses were conducted in R 4.2.2 (58). Model diagnostics were checked using simulated residuals with *DHARMa* (v 0.4.6) (59) and significance of fixed or random effects were assessed with likelihood ratio tests. Transformations and distributions for each modeled response variable are as follows: *Shoot mass*, *nodule width*, and *CFU/nodule* were log-transformed to improve fit to gaussian error distributions implemented with *lme4* (v 1.1-35.3) (60). For *transconjugants per mating*, *number of nodules*, and *number of N-fixing* (*pink*) *nodules*, we used the negative binomial distribution in models implemented with *glmmTMB* (v 1.1.9) (61). For *%N* in leaf tissue, we expressed the percent as a proportion and used the *glmmTMB* beta family distribution. For *%N* in leaf tissue models, we also included a dispersion parameter representing a binary factor for nodule color (*pink* or *white*) to improve heteroskedasticity due to differences in error dispersion between plants with only white nodules vs plants with one or more pink nodules.

### Broad sense heritability

We first quantified broad sense heritability (*H^2^*) for SI *transmission*. We partitioned the effect of the donor-SI genotype (*V_D_*), recipient genotype (*V_R_*), and their interaction (*V_DxR_*) on the number of *transconjugants per mating* (*V_P_*) across all 180 mating trials (62,63) using a random effects model. Since we cannot parse the variance attributed to the donor chromosome from the donor SI, we refer to genotypic effects of SI+ donor genotypes as ‘donor-SI’.

We next quantified the impact of the donor-SI vs recipient genotype on *symbiotic function* of novel transconjugants. We partitioned the impact of donor-SI genotype (*V_D_*), recipient genotype (*V_R_*), and experimental block (*V_E_*) on plant and rhizobium symbiotic fitness (*V_P_*) for transconjugant-inoculated plants in the greenhouse experiment using random effects models.

We standardized sources of variance by dividing them by the total variance of the trait analyzed. Variance components for *transconjugants per mating*, *%N*, *nodule number* and *Nfixing nodule number* were standardized to the total model explained variance since residual variance was not calculated for non-gaussian models (47). We obtained 95% confidence intervals for each variance component using the percentile method for 10,000 nonparametric bootstrap replicates of each model (*boot* v 3.13-30) (64). *CFU/nodule* was a response variable in this variance component model and not in the models described below because of small sample size (see Supplemental Information).

### Symbiotic fitness (donor vs. transconjugant models)

To determine the extent to which SI function depends upon the chromosome genotype in which it resides, we compared endosymbiosis outcomes for the SI+ donor vs transconjugants that received the donor’s SI. We used generalized linear mixed models (GLMMs) to model symbiosis response variables with rhizobium genotype as a fixed effect, and block as a random effect. For each symbiosis trait, we assessed the significance of rhizobium genotype using a likelihood ratio test followed by estimated marginal means (*emmeans* v 1.10.1) (65) post hoc analyses among genotypes. Uninoculated negative controls were included in the *shoot mass* and *%N* models to assess which strains conferred benefit to the host plant and significance was unaltered by inclusion of this data.

### Eco-evolutionary predictors (eco-evo models)

We tested whether ecological or evolutionary (*eco-evo*) attributes of donor-SI-recipient mating pairs predicted SI transmission or function. Eco-evo attributes were recorded during original strain isolation, including geographic coordinates, host plant species (*A. wrangelianus* or *A. brachycarpus),* and soil type (serpentine or non-serpentine) (41). We calculated geographic and phylogenetic pairwise distances between each donor-SI and recipient strain, and binary variables represented whether the donor-SI and recipient originated from the same or different host plant species or soil type.

We modeled the number of *transconjugants per mating*, host plant fitness, or rhizobium fitness as response variables in GLMMs with phylogenetic distance, geographic distance, host plant species and soil type as fixed effects, and experimental block (except not for SI transfer), donor-SI, recipient, and their interaction as random effects. When all random effects were included in models, *%N* and *N-fixing nodule number eco-evo models* were overparametrized. Therefore, we removed the donor-SI or interaction random effect variance component, respectively, because they were estimated to be nearly zero in these models (66).

## RESULTS

Symbiosis island transfer was found to be common, occurring in 80% (144/180) of mating pairs across this wild *Mesorhizobium* population (Figure 2a). We confirmed SI presence and the expected recipient genotype via PCR and sequencing for 15 of the 144 successful matings (Table S2; Table S3; Figure 3). Confirming expected replicon identity in all cases, ANI between wild-type donor and SI-marked donor, and between transconjugants and respective recipients, is nearly 99.9%, while other pairwise ANI are less than 99% (Table S4).

**Figure 2.**
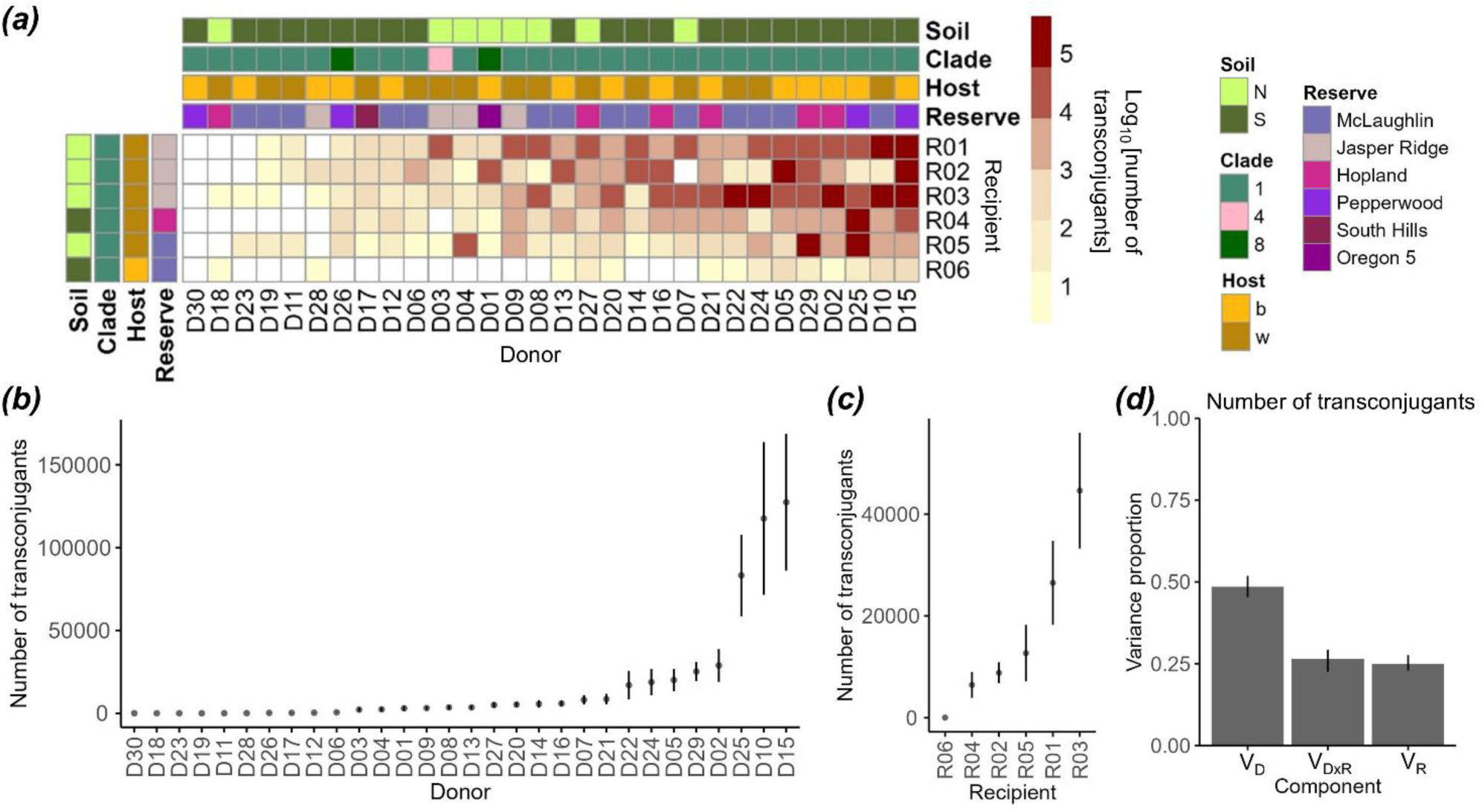
**Genetic determinants of symbiosis island (SI) transfer**. **(a)** Heatmap of the mean Log_10_ number of transconjugants for each pairwise mating (n =180) from high (dark red) to low (beige) transfer. White boxes indicate no SI transfer was detected. Additional colored boxes show donor and recipient eco-evo factors. Soil type: Serpentine soil (S), non-serpentine soil (N). Clade: clade designation based on 354 conserved marker genes. Host: *Acmispon brachycarpus* (b), *A. wrangelianus* (w). Reserve: The reserve in Oregon or California where the strains were collected. The mean number of transconjugants for each **(b)** donor and **(c)** recipient. Filled circles show genotype mean values +/- standard error. **(d)** The proportion of phenotypic variance in the number of transconjugants per mating explained by donor-SI (*V_D_*), recipient (*V_R_*), and their interaction (*V_DxR_*). Bars show variance estimates and 95% confidence intervals.

**Figure 3.**
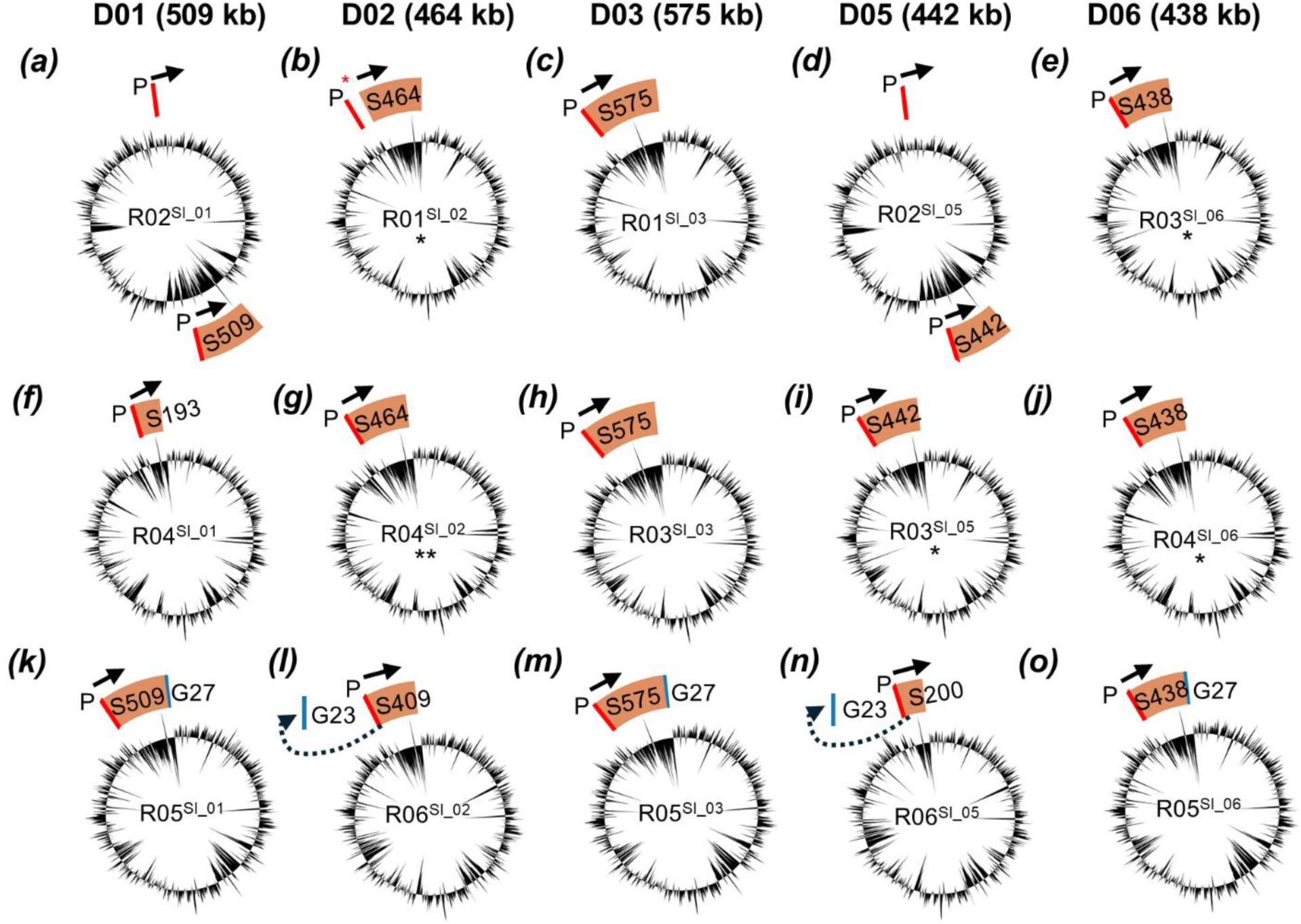
SI gain transconjugant genomes reveal complete and partial SI transfer. Donor-SI IDs and size are shown above three transconjugant genomes that gained the respective SI **(a-o)**, each with its origin of replication closest to the top of the page. Shown is the SI insertion site at a Phe-tRNA gene (P; red and enlarged for visualization) in the 5’ to 3’ direction (black arrow) such that the 17-bp direct repeat of the 3’ end of the Phe-tRNA gene is at the end of the SI (S; pink). Transconjugants are classified as mutualists if they fix nitrogen (asterisk). Only one transconjugant conferred a significant increase in shoot mass (two asterisks). The fate of inferred genomic islands (G; blue) originally inserted at the recipient Phe-tRNA gene is shown: in **l** and **n** these were replaced (dotted arrow) whereas in **m** and **o** these were moved downstream due to SI insertion. The number after S or G indicates size (kb). Genomic maps were made in Proksee (52).GC content is shown in black. In **b**, low sequencing coverage is indicated with a red asterisk (Table S3).

All sequenced transconjugants contained sequence from the expected donor-SI at the Phe-tRNA gene (Figure 3). However, for 3/15 strains only a partial fraction of the donor-SI was inserted: transconjugants R04^SI_01^, R06^SI_02^, and R06^SI_05^ are missing 62%, 12%, or 55% of the SI, respectively (Figure 3). 5/15 of the transconjugants were derived from two recipient strains that had putative GIs originally inserted at the Phe-tRNA gene. For two of these transconjugants, the GI was replaced by the donor-SI (R06^SI_02^ and R06^SI_05^). For three of these transconjugants, the GI was retained directly downstream of the donor-SI (R05^SI_01^, R05^SI_03^, and R05^SI_06^; Table S3; Figure 3).

The SI acquisition readily conferred endosymbiosis to recipient strains, which resulted in symbiotic fitness benefits to rhizobia, though benefit to the host plant was less common. SI acquisition enabled all tested transconjugants to form nodules, transforming previously nonnodulating strains into nodulating endosymbionts. In plating assays, we detected 2.6 × 10^5^ CFU per nodule on average for 13/15 of the transconjugants we tested, demonstrating that these novel endosymbionts can proliferate within nodules and gain symbiotic fitness (progeny from R03^SI_03^ and R06^SI_05^ could not be recovered from nodules). However, donors formed greater CFU per nodule on average (6.1 × 10^6^) compared to the transconjugants. Furthermore, the benefits gained from symbiosis differed among donor-SI and recipient genotypic combinations. We observed epistasis whereby 5/15 transconjugants bearing a particular SI formed fewer and smaller nodules compared to the donor that also carries this SI (e.g., D01^SI_01^ and D03^SI_03^; Figure 4f,h). On the other hand, 5/15 transconjugants bearing a particular SI formed nodules that were indistinguishable in size and number compared to the donor that also carries this SI (e.g., D02^SI_02^ and D06^SI_06^; Figure 4f,h).

**Figure 4:**
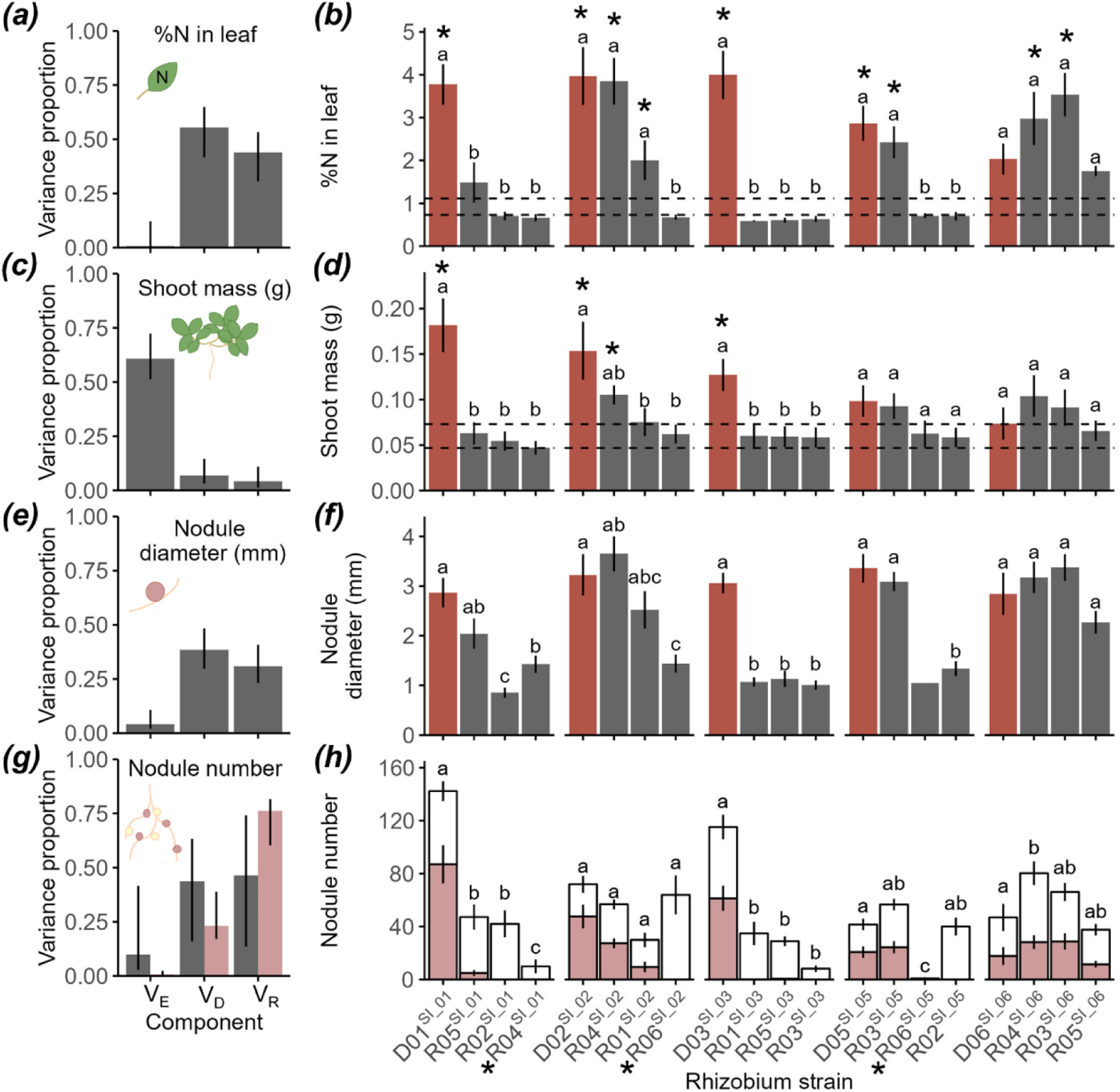
Symbiosis island acquisition generates novel nodulating commensal or mutualistic endosymbionts, depending on donor and recipient genotype and their GxG interaction. The percent of phenotypic variance in the **(a)** percent of nitrogen in leaf tissue, **(c)** shoot mass, and rhizobium **(e)** nodule width and **(g)** nodule number (grey bars) or N-fixing nodule number (pink bars) explained by experimental block (V_E_), donor-SI genotype (V_D_) and recipient genotype (V_R_). Bars show variance estimates and 95% confidence intervals. **(b)** The percent of nitrogen in leaf tissue, **(d)** shoot mass, **(f)** nodule width and **(h)** nodule number for SI donors and SI gain transconjugants. Within each donor SI type (SI_01, SI_02, SI_03, SI_05, and SI_06), the donor is shown first (D01, D02, D03, D05, and D06), followed by the three transconjugants (named by recipient genotype R01 - R06). Donors are colored red in the shoot mass and nodule width plots, while nodule number plot colors show the proportion of pink and white nodules. Bars are genotypic means +/- standard error. Letters above bars indicate significant differences in phenotypes among the donor and transconjugants within each SI type and asterisks indicate differences from the rhizobia-free controls at *P* < 0.05. An asterisk by the rhizobium strain ID indicates the strain contains a partial SI.

For a third of transconjugants, SI acquisition transformed non-nodulating strains into endosymbionts able to fix nitrogen within nodules on a host plant. Consistent with functional symbiotic nitrogen fixation, 4/5 of the SI donor strains and 5/15 transconjugants resulted in *%N* greater than that of uninoculated control plants (Figure 4b). For one transconjugant, SI acquisition also enhanced shoot growth. Consistent with a net fitness benefit to the plant, 3/5 donors and 1/15 transconjugants increased shoot mass compared to uninoculated controls (*donor vs transconjugant models;* Figure 4d).

### Broad sense heritability of SI transmission and function

Donor-SI genotype (*V_D_*), the recipient genotype (*V_R_*), and their interaction (*V_DxR_*) all contributed to broad sense heritability of SI transfer (Table S5; Figure 2). For number of *transconjugants per mating*, 48.5% of the variation is explained by donor-SI genotype (*V_D_*), 25% is explained by recipient genotype (*V_R_*), and 26.5% is explained by a donor-recipient genotype interaction (*V_DxR_*) (Table S5; Figure 2d). Bootstrap confidence intervals indicate variance explained by *V_D_* is greater than both *V_R_* and *V_DxR_*, while *V_R_* and *V_DxR_* confidence intervals overlap (Table S5; Figure 2d).

Both the donor-SI and recipient genotype predict the functional consequences of SI transmission for rhizobia and their plant hosts. For plant *shoot mass*, donor-SI and recipient genotype explain 6.9% and 4.1% of the total variance, respectively, while environmental variation in the greenhouse (block), explained 60.8% of the variation (Table S6; Figure 4c). In contrast, other phenotypes were remarkably insensitive to environmental variation in the greenhouse. For *%N* in leaf tissue, donor-SI explained 55.5% of the variance, and recipient genotype explained 43.9%, and experimental block was negligible (Table S6; Figure 4a). For rhizobium fitness, variance explained by block was less than 5% across all measures and was not significant in models for the *number of nodules* or *CFU/nodule* (Table S6; Figure 4e,g; Figure S4a). Donor-SI and recipient genotype explained 38.5% and 31.1% of the variance in *nodule width* (Table S6; Figure 4e) and 33% and 19% of the variance in *CFU/nodule*, respectively (Table S6; Figure S2). *Nodule width* and *CFU/nodule* were positively correlated (Figure S3). For the *number of nodules* and *number of N-fixing nodules*, donor-SI genotype explained 43.7% or 23.1% of the variance, while recipient genotype explained 46.4% or 76.2% of the variance, respectively (Table S6; Figure 4g). Donor-SI genotype explained indistinguishable amounts of the variance for all rhizobium phenotypes (*nodule width*, *CFU/nodule*, *nodule number, N-fixing nodule number*), while recipient genotype explained more variance in *N-fixing nodule number* compared to other rhizobium phenotypes.

### Evolutionary factors predict SI transmission fitness effects

Phylogenetic and ecotypic factors were associated with the degree to which SI transmission altered symbiosis function. Greater phylogenetic relatedness between the SI donor and recipient led to transconjugants with greater symbiotic nitrogen fixation (*%N*: *χ*^2^ = 14.6, *P* < 0.001; Figure 5a), *shoot mass* (*χ*^2^ = 7.9, *P* = 0.005; Figure 5b), *nodule width* (*χ*^2^ = 16.2, *P* < 0.001; Figure 5c), and *N-fixing nodule number* (*χ*^2^ = 19.9, *P* < 0.001; Figure 5d) (Table S7). In addition, SI donors and recipients that originated from the same soil type, either serpentine or non-serpentine, resulted in transconjugants that conferred greater *%N* (*χ*^2^ = 6.4, *P* = 0.011; Figure 5e), *shoot mass* (*χ*^2^ = 5.6, *P* = 0.018; Figure 5f), *nodule width* (*χ*^2^ = 5.1, *P* = 0.024; Figure 5g), and *N-fixing nodule number* (*χ*^2^ = 19.3, *P* < 0.001; Figure 5h) (Table S7). No eco-evo factors predicted *nodule number* (Table S7) or *transconjugants per mating* (Table S8).

**Figure 5.**
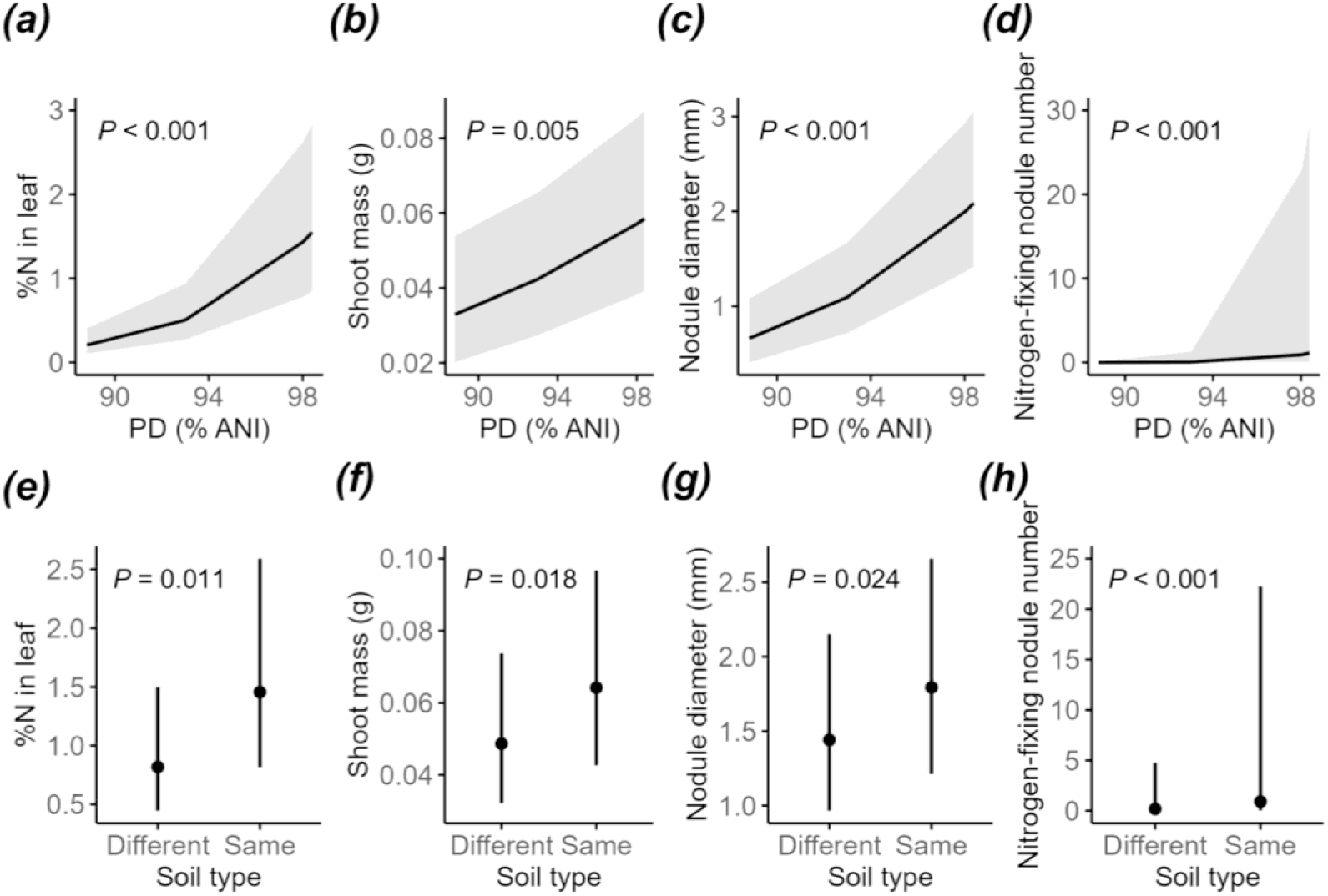
Eco-evo factors predict SI function. Eco-evo factors also predict SI function in transconjugants following acquisition, where greater phylogenetic relatedness is positively correlated with (a) the percent of nitrogen in leaf tissue, (b) shoot mass, (c) nodule width, and (d) the number of nitrogen-fixing nodules. Black line, marginal effects; grey shading, 95% confidence interval. (e) The percent of nitrogen in leaf tissue, (f) shoot mass, (g) nodule width, and (h) nitrogen-fixing nodule number are also higher when SI donors and recipients came from the same soil type compared to donors and recipients from different soil types. Filled circles, marginal effects with 95% confidence intervals. Significance values in the prediction plots are from likelihood ratio tests.

## DISCUSSION

Our experiments provide genetic, genomic, and functional evidence that mutualistic endosymbiosis can be an evolutionarily labile trait driven by segregating presence/absence variation for an MGE in wild rhizobia. To overcome the limitations of using contemporary interactions to infer critical events during the onset of endosymbiosis (1), we generated novel endosymbioses by transferring rhizobia symbiosis islands (SIs) into non-nodulating strains that lacked the SI. SI transmission was sufficient to generate *de novo* root nodule endosymbioses for wild SI donor and recipient strains. Most novel endosymbionts originated as commensals that incurred no detectable cost or benefit to the host in clonal inoculation, in contrast to predictions that novel symbionts tend to exploit a host (1,10,11). In fact, a third of novel endosymbionts originated as *de novo* mutualists that provided symbiotically fixed nitrogen to the host, consistent with broad genetic potential in non-nodulating strains to benefit a plant after a single MGE acquisition. We thus elucidate how MGEs can potentiate and constrain major evolutionary transitions to expand bacterial niches, with cascading effects on host organisms (20,67).

We find abundant heritable variation for the propensity for SI transmission and function among donor and recipient genotypes, and their GxG interactions. Thus, the population could evolve in response to selection on transmission to increase or decrease the propensity for niche expansion into endosymbiosis. Consistent with phylogenetic limits to transfer of MGE function, novel endosymbionts derived from more closely related SI donor and recipient strains showed greater nitrogen fixation. However, we did not detect phylogenetic limits to SI HGT, which could reflect selfish selection for generalized HGT of this mobile element (22,26,68). In fact, in multiple instances, the SI was able to displace other genomic elements residing at its characteristic tRNA gene insertion site. Rarely, a large section of the SI was lost at insertion, generating strains capable of nodulation but not nitrogen fixation, which are unlikely to evolve into mutualists due to extensive gene loss.

### Incipient endosymbiosis

The non-nodulating kin of nodulating rhizobia, are widespread and diverse in plant, soil, and even animal gut microbiomes (69). However, little is known about their evolutionary dynamics. We demonstrate experimentally that co-occurring nodulating rhizobia and their nonnodulating kin are commonly able to transmit an MGE conferring mutualistic functions such that endosymbiosis is evolutionarily labile and driven by segregating presence/absence variation among wild *Mesorhizobium* from across Western USA. Conjugative transfer of the SI was relatively common evolutionary outcome for most of the matings we attempted. Here, SI transmission was successful in 80% of the 180 pairwise matings between 30 SI+ donors and 6 SI- recipients. However, as others have found, SI transfer was relatively infrequent among individual cells in a mating, producing an average of 1.7 × 10^4^ transconjugants per mating spot (24,36,48).

The first step during a transition to endosymbiosis is often hypothesized to arise via exploitation due to selection for one partner to benefit at the other’s expense (1,11). However, we find that a symbiotic fitness benefit for a novel endosymbiont is common and often incurs little cost to the host, despite host investment in symbiotic structures like nodules (24,36,37). While all transconjugants we tested induce root nodule formation and thus gained a symbiosis fitness benefit, none incurred measurable harm to the host, consistent with little role for exploitation during the transition from non-nodulating to nodulating endosymbiotic rhizobia. Instead, two thirds of novel root nodule endosymbionts we tested originated as commensals that incurred no major cost or benefit to the host. Thus, we infer that a commensal symbiont lifestyle may often underpin early steps along the transition to endosymbiosis via symbiosis MGE acquisition, in interactions where the costs of initiating endosymbiosis for hosts are low. Here, selection on the host to prevent infection by suboptimal novel symbionts may be weak or ineffective, especially if the host can discriminate against suboptimal symbionts at later time points in the interaction via sanctions (70) maintaining a niche for novel endosymbionts during initiation of endosymbiotic infection (12).

*De novo* mutualism was also a common first step during incipient endosymbiosis: a third of the tested strains that gained the SI were capable of both benefitting from, and conferring benefit to, the legume host. Incipient endosymbiosis is not typically hypothesized to evolve via initial mutualism because the acquisition of mutations or loci conferring endosymbiosis would have to simultaneously benefit the chromosome bearing them as well as the host, which is a restrictive set of conditions (1). In our study, the relatively close phylogenetic relatedness between donor and recipient strains may result in pre-adaptation or co-evolution between the recipient lineages and the SI, such that an MGE that can confer beneficial endosymbiosis continues to do so in novel genomic contexts. This outcome may reflect past selection for generalist SI genotypes that obtain high fitness via HGT because they can benefit diverse host lineages, unlike specialist SI genotypes that are more restricted to vertical transmission with the host genome because they are beneficial to few host lineages.

### Genetic variation in SI transmission

In the *Mesorhizobium* we examine, the propensity to transfer or acquire the SI, and the symbiotic traits it confers, are genetically variable traits that could shift in response to selection. While several studies find variation in conjugative transfer rates among individual donor and recipient lineages (27,36,71), little is known about how donor genotype, recipient genotype, or their GxG interaction affect transmission in populations. In our trials, the propensity to transmit the SI was highly genetically determined: broad sense heritable variation among donor-SI genotypes explained nearly half (49%) of variation in transmission. This was greater than the influence of the recipient genotype, and the donor-SI-recipient GxG interaction, which each explained a quarter of variation in SI transmission rates (25% and 27%, respectively). Thus, donors, and to a lesser extent, recipients, bear loci that have consistent genetic impacts on transmission rates. Some loci impacting transmission also appear subject to epistasis, such that their impact on transmission is sensitive to the identity of an interacting genotype. This contrasts with findings in other systems that show a weak role for recipient genotype in determining the outcome of MGE conjugative transfer (72,73). For example, Allard et al. (2023) implicated few chromosomal genes in variation in conjugation frequency despite screening ∼4000 *E. coli* gene deletion mutants as recipients. However, such studies often test dispensable genes in recipients, whereas recipient genes that influence conjugation could have large effects on survival. In natural populations of rhizobia, SI acquisition by non-nodulating recipients may represent a favorable lifestyle switch when host legumes are abundant, and chromosomes bear variants that promote symbiotic function.

Genetic variation for SI transmission could be maintained by ongoing coevolution between the SI and the main chromosome. Antagonistic coevolution could occur because while the SI gains fitness from HGT among chromosomal lineages, this comes at a cost to the chromosome it resides in due to the metabolic and fitness costs of conjugative transfer (68,74,75). Thus, while SIs can evolve to carry loci that promote their conjugative transfer and HGT (76–79), chromosomes can reciprocally accumulate loci that inhibit SI transmission (35).

However, antagonistic coevolution could be tempered under conditions where selection to maintain investment in the shared cell they reside in and vertical transmission to daughter cells is sufficient to attenuate selection on the SI for costly conjugation (22,23,36,74). Future studies are needed to identify loci that modulate SI transfer rates in rhizobia and test these predictions regarding whether they reside in the SI or chromosome.

Beyond coevolution, genetic variation for SI transmission could also be maintained by fluctuating selection due to variable host plant distributions. The SI confers symbiotic fitness benefits where compatible host plants occur (74,75). Thus, proximity to host legumes could favor loci that enhance chromosomal SI gain and SI function, enhancing the fitness of chromosomal lineages, such as R01, that are able to acquire the SI from many donors (24,38). Conversely, in the absence of compatible host plants, selection could favor chromosomal resistance to acquiring the SI. The SI comprises 8% of the genome and in the absence of compatible host plants, the cost of conjugation and replicating the SI may outweigh any benefit (80,81). Bacterial resistance to MGEs is common via mechanisms such as cell surface receptor compatibility for conjugation, or CRISPR-Cas and restriction modification systems (35,72,82,83). Where compatible hosts plants are rare, chromosomal lineages, such as R06, for which SI transmission was only successful with half of the donors, may benefit from resisting SI transmission.

### Genetic variation in SI function

SI donor and recipient *Mesorhizobium* strains differ in their main genotypic effects on symbiosis traits, and also show epistatic genotypic effects such that their impact on symbiosis traits is sensitive to the identify of an interacting genotypes, consistent with findings from other rhizobia (24,36,84,85) Multiple SIs conferred higher shoot mass and nitrogen fixation in their native, co-adapted chromosomal context than within a novel chromosomal background, for example, D01 and D03. In one case, an SI (D06) conferred nitrogen fixation in two recipient chromosomal genotypes, despite conferring no significant fixation in its native context. However, in other cases SIs provided indistinguishable shoot mass benefits in native and novel chromosomal backgrounds (D02 and D05). These findings are consistent with variation in recipient chromosome pre-adaptations that influence symbiotic function upon uptake of a particular SI. This variation could reflect a history of variable selection on symbiotic function, even in lineages that now lack the SI, or could reflect other processes such as indirect selection on traits linked to symbiotic function (38,74,75). Rhizobium symbiotic fitness phenotypes, such as nodule width and N-fixing nodule number, showed similarly strong genetic variation and epistatic effects.

Closer phylogenetic relatedness between donor and recipient *Mesorhizobium* led to greater symbiotic function in resulting transconjugants. Closely related donor-recipient matings generated transconjugants that conferred higher plant shoot mass, nitrogen fixation, nodule width, and number of N-fixing nodules. For example, transconjugants with an SI from D03, a distantly related clade 4 donor, formed small white nodules and did not fix nitrogen when acquired by clade 1 recipients. However, when these recipients received an SI from a donor in their same clade (D02, D05, and D06), they formed large pink, nitrogen fixing nodules. Such limitations to SI function in recipient backgrounds may hinge on co-adaptation of the donor-SI and recipient over evolutionary time (38,45). In other rhizobia, when symMGEs are transferred among genera or to non-rhizobia, transconjugants are unable to fix nitrogen or compete with the native donor-SI strain for nodulation, and the symMGE is susceptible to loss (86–88). While MGEs are modular genetic elements that bear a complete suite of loci to enact particular cellular functions, phylogenetic divergence among donor and recipient lineages may limit MGE function across diverse microbial taxa.

Matings between donors and recipients from the same soil ecotype tended to generate transconjugants with greater SI function, consistent with epistasis between the loci conferring environmental and symbiotic adaptation. The *Mesorhizobium* in our study have repeatedly evolved two soil ecotypes. Serpentine soil ecotypes adapt to toxic nickel in serpentine soils via acquisition of the nickel resistance island (NRI), that often occurs as an integrative and conjugative element (ICE) (41). Non-serpentine soil ecotypes occur elsewhere, are sensitive to nickel, and usually lack the NRI (41). In our experiment, donor-recipient pairings of the same soil ecotype generated transconjugants that conferred greater plant shoot mass, nitrogen fixation, nodule width, and number of N-fixing nodules than transconjugants from donor-recipient pairings from different soil ecotypes. Thus, the SI may have adapted to function optimally in either the presence or absence of the NRI within the cellular replicon. These findings are consistent with a scenario in which large adaptive ICEs impact each other’s function and lead to intragenomic co-adaptation (17). Here, moving the SI into a novel genomic context, either the novel presence or absence of the NRI, could then lead to reduced symbiosis function. Thus, ecotypic adaptation to environmental variation appears to incur genetic constraints on symbiotic function, highlighting an evolutionary tradeoff between MGE-conferred adaptation to environment vs host (1).

While we observed phylogenetic and ecotypic patterns for SI transmission and function, our findings rest upon a relatively limited sample size of transconjugants. Larger functional studies are needed to more fully reveal the complex evolutionary interplay of adaptive MGEs in the bacterial mobilome. Other evolutionary attributes of donor and recipients we tested, host plant species or geographic distance between donor and recipient collection sites were not associated with SI transmission or function. Further studies could more robustly test for multiple ecological and evolutionary factors to predict SI transmission and function.

## Conclusion

To harness the adaptive traits MGE confer for human benefit, such as producing superior agricultural inoculants or probiotics, it’s critical to understand the interplay between donors, recipients and MGEs (25,27,89). We reveal that the propensity to transfer or acquire the SI is a heritable trait that could respond to selection to alter transmission rates, and thus, contribute to the evolvability of *Mesorhizobium* lineages. The functional consequences of gaining the SI are also heritable traits. The ability to form nodules in novel endosymbioses appears to simply be driven by acquiring the SI, while the ability to fix nitrogen and confer mutualistic benefits depends on the recipient background genome. While the transconjugants we examined confer benefit to the host plant in one-third of cases, it is possible that if we evolved transconjugants on host plants they could become more beneficial, as MGEs and recipient chromosomes can coadapt over time (27).

## DATA AVAILABILITY

Genome sequence data will be deposited in NCBI and data will be deposited in the Dryad data repository, but is currently private for peer review.

## ACKNOWLEDGEMENTS

SSP and APM were supported by US National Science Foundation grant DEB-1943239 to SSP. APM was also supported by the American Association of University Women (AAUW) American Dissertation Fellowship and the Washington State University, School of Biological Sciences, Rexford Daubenmire Fund for Graduate Education. This research used resources of the Center for Institutional Research Computing at Washington State University.

## Notes

### Competing Interest Statement

The authors have declared no competing interest.

## REFERENCES

1. Brockhurst MA, Cameron DD, Beckerman AP. Fitness trade-offs and the origins of endosymbiosis. PLOS Biology [Internet]. 2024 Apr 12 [cited 2025 May 16];22(4):e3002580. Available from: https://journals.plos.org/plosbiology/article?id=10.1371/journal.pbio.3002580

2. Richards TA, Moran NA. Symbiosis: In search of a deeper understanding. PLOS Biology. 2024 Apr 12;22(4):e3002595.

3. Vosseberg J, van Hooff JJE, Köstlbacher S, Panagiotou K, Tamarit D, Ettema TJG. The emerging view on the origin and early evolution of eukaryotic cells. Nature. 2024 Sept;633(8029):295–305.

4. Bordenstein SR, The Holobiont Biology Network. The disciplinary matrix of holobiont biology. Science [Internet]. 2024 Nov 15 [cited 2025 Sept 9];386(6723):731–2. Available from: https://www.science.org/doi/10.1126/science.ado2152

5. McCutcheon JP. The Genomics and Cell Biology of Host-Beneficial Intracellular Infections. Annual Review of Cell and Developmental Biology [Internet]. 2021 Oct 6 [cited 2025 Sept 9];37(Volume 37, 2021):115–42. Available from: https://www.annualreviews.org/content/journals/10.1146/annurev-cellbio-120219-024122

6. Bock R. Witnessing Genome Evolution: Experimental Reconstruction of Endosymbiotic and Horizontal Gene Transfer. Annual Review of Genetics [Internet]. 2017 Nov 27 [cited 2025 Sept 9];51(Volume 51, 2017):1–22. Available from: https://www.annualreviews.org/content/journals/10.1146/annurev-genet-120215-035329

7. Puri KM, Butardo V, Sumer H. Evaluation of natural endosymbiosis for progress towards artificial endosymbiosis. Symbiosis [Internet]. 2021 May 1 [cited 2025 Sept 9];84(1):1–17. Available from: 10.1007/s13199-020-00741-5

8. Frank SA. Models of Symbiosis. The American Naturalist [Internet]. 1997 July [cited 2025 Sept 10];150(S1):S80–99. Available from: https://www.journals.uchicago.edu/doi/10.1086/286051

9. Keeling PJ, McCutcheon JP. Endosymbiosis: The feeling is not mutual. Journal of Theoretical Biology [Internet]. 2017 Dec 7 [cited 2025 Sept 9];434:75–9. Available from: https://www.sciencedirect.com/science/article/pii/S0022519317302795

10. Law R, Dieckmann U. Symbiosis through exploitation and the merger of lineages in evolution. Proc Biol Sci [Internet]. 1998 July 7 [cited 2025 Sept 10];265(1402):1245–53. Available from: https://www.ncbi.nlm.nih.gov/pmc/articles/PMC1689185/

11. Sachs JL, Skophammer RG, Regus JU. Evolutionary transitions in bacterial symbiosis. Proceedings of the National Academy of Sciences. 2011 June 28;108(Supplement_2):10800–7.

12. Porter SS, Dupin SE, Denison RF, Kiers ET, Sachs JL. Host-imposed control mechanisms in legume–rhizobia symbiosis. Nat Microbiol [Internet]. 2024 Aug 2 [cited 2024 Aug 3];1–11. Available from: https://www.nature.com/articles/s41564-024-01762-2

13. Wright RCT, Wood AJ, Bottery MJ, Muddiman KJ, Paterson S, Harrison E, et al. A chromosomal mutation is superior to a plasmid-encoded mutation for plasmid fitness cost compensation. PLOS Biology [Internet]. 2024 Dec 2 [cited 2025 Feb 25];22(12):e3002926. Available from: https://journals.plos.org/plosbiology/article?id=10.1371/journal.pbio.3002926

14. Azuma Y, Tsuru S, Habuchi M, Takami R, Takano S, Yamamoto K, et al. Synthetic symbiosis between a cyanobacterium and a ciliate toward novel chloroplast-like endosymbiosis. Sci Rep [Internet]. 2023 Apr 13 [cited 2025 Sept 9];13(1):6104. Available from: https://www.nature.com/articles/s41598-023-33321-w

15. Cournoyer J, Altman SD, Gao Y le, Wallace CL, Zhang D, Lo GH, et al. Engineering artificial photosynthetic life-forms through endosymbiosis. Nat Commun [Internet]. 2022 Apr 26 [cited 2025 Sept 9];13(1):2254. Available from: https://www.nature.com/articles/s41467-02229961-7

16. Giger GH, Ernst C, Richter I, Gassler T, Field CM, Sintsova A, et al. Inducing novel endosymbioses by implanting bacteria in fungi. Nature [Internet]. 2024 Oct 2 [cited 2024 Oct 4];1–8. Available from: https://www.nature.com/articles/s41586-024-08010-x

17. Lang AS, Buchan A, Burrus V. Interactions and evolutionary relationships among bacterial mobile genetic elements. Nat Rev Microbiol [Internet]. 2025 Mar 11 [cited 2025 Apr 13];1–16. Available from: https://www.nature.com/articles/s41579-025-01157-y

18. Arnold BJ, Huang IT, Hanage WP. Horizontal gene transfer and adaptive evolution in bacteria. Nat Rev Microbiol. 2022 Apr;20(4):206–18.

19. Pena MM, Bhandari R, Bowers RM, Weis K, Newberry E, Wagner N, et al. Genetic and Functional Diversity Help Explain Pathogenic, Weakly Pathogenic, and Commensal Lifestyles in the Genus Xanthomonas. Genome Biology and Evolution [Internet]. 2024 Apr 1 [cited 2025 Apr 13];16(4):evae074. Available from: 10.1093/gbe/evae074

20. Porter S, Holtappels D, Montoya A, Koskella B. Causes and consequences of bacterial local adaptation via MGEs in the plant microbiome. New Phytologist [Internet]. 2025 [cited 2025 Dec 1]; Available from: https://onlinelibrary.wiley.com/doi/abs/10.1111/nph.70766

21. Weisberg AJ, Rahman A, Backus D, Tyavanagimatt P, Chang JH, Sachs JL. Pangenome Evolution Reconciles Robustness and Instability of Rhizobial Symbiosis. mBio [Internet]. 2022 Apr 13 [cited 2025 Apr 3];13(3):e00074-22. Available from: https://journals.asm.org/doi/full/10.1128/mbio.00074-22

22. Brockhurst MA, Harrison E, Hall JPJ, Richards T, McNally A, MacLean C. The Ecology and Evolution of Pangenomes. Current Biology. 2019 Oct 21;29(20):R1094–103.

23. Virolle C, Goldlust K, Djermoun S, Bigot S, Lesterlin C. Plasmid Transfer by Conjugation in Gram-Negative Bacteria: From the Cellular to the Community Level. Genes. 2020 Nov;11(11):1239.

24. Colombi E, Hill Y, Lines R, Sullivan JT, Kohlmeier MG, Christophersen CT, et al. Population genomics of Australian indigenous Mesorhizobium reveals diverse nonsymbiotic genospecies capable of nitrogen-fixing symbioses following horizontal gene transfer. Microbial Genomics [Internet]. 2023 Jan 5 [cited 2023 Jan 12];9(1). Available from: https://www.microbiologyresearch.org/content/journal/mgen/10.1099/mgen.0.000918

25. Bottery M, Brockhurst MA. Rapid evolution helps bacteria to pick up a plasmid. Proceedings of the National Academy of Sciences. 2023 May 9;120(19):e2304474120.

26. Koonin EV. Horizontal gene transfer: essentiality and evolvability in prokaryotes, and roles in evolutionary transitions. F1000Research. 2016 July 25;5:F1000 Faculty Rev.

27. Benz F, Hall AR. Host-specific plasmid evolution explains the variable spread of clinical antibiotic-resistance plasmids. Proceedings of the National Academy of Sciences. 2023 Apr 11;120(15):e2212147120.

28. Brockhurst MA, Harrison E. Ecological and evolutionary solutions to the plasmid paradox. Trends in Microbiology. 2022 June 1;30(6):534–43.

29. Alderliesten JB, Duxbury SJN, Zwart MP, de Visser JAGM, Stegeman A, Fischer EAJ. Effect of donor-recipient relatedness on the plasmid conjugation frequency: a meta-analysis. BMC Microbiol. 2020 May 26;20(1):135.

30. Chu BTT, Petrovich ML, Chaudhary A, Wright D, Murphy B, Wells G, et al. Metagenomics Reveals the Impact of Wastewater Treatment Plants on the Dispersal of Microorganisms and Genes in Aquatic Sediments. Appl Environ Microbiol [Internet]. 2018 Mar 1 [cited 2021 Apr 20];84(5). Available from: https://aem.asm.org/content/84/5/e02168-17

31. Greenlon A, Chang PL, Damtew ZM, Muleta A, Carrasquilla-Garcia N, Kim D, et al. Global-level population genomics reveals differential effects of geography and phylogeny on horizontal gene transfer in soil bacteria. PNAS. 2019 July 23;116(30):15200–9.

32. Hanson CA, Fuhrman JA, Horner-Devine MC, Martiny JBH. Beyond biogeographic patterns: processes shaping the microbial landscape. Nature Reviews Microbiology. 2012 July;10(7):497–506.

33. Martiny JBH, Bohannan BJM, Brown JH, Colwell RK, Fuhrman JA, Green JL, et al. Microbial biogeography: putting microorganisms on the map. Nature Reviews Microbiology. 2006 Feb;4(2):102–12.

34. Dimitriu T. Evolution of horizontal transmission in antimicrobial resistance plasmids. Microbiology. 2022;168(7):001214.

35. Horne T, Orr VT, Hall JP. How do interactions between mobile genetic elements affect horizontal gene transfer? Current Opinion in Microbiology. 2023 June 1;73:102282.

36. Haskett TL, Terpolilli JJ, Bekuma A, O’Hara GW, Sullivan JT, Wang P, et al. Assembly and transfer of tripartite integrative and conjugative genetic elements. Proc Natl Acad Sci U S A. 2016 Oct 25;113(43):12268–73.

37. Hill Y, Colombi E, Bonello E, Haskett T, Ramsay J, O’Hara G, et al. Evolution of Diverse Effective N2-Fixing Microsymbionts of Cicer arietinum following Horizontal Transfer of the Mesorhizobium ciceri CC1192 Symbiosis Integrative and Conjugative Element. Applied and Environmental Microbiology. 2021 Feb 12;87(5):e02558–20.

38. Liu S, Jiao J, Tian CF. Adaptive Evolution of Rhizobial Symbiosis beyond Horizontal Gene Transfer: From Genome Innovation to Regulation Reconstruction. Genes. 2023 Feb;14(2):274.

39. Nandasena KG, O’Hara GW, Tiwari RP, Sezmiş E, Howieson JG. In situ lateral transfer of symbiosis islands results in rapid evolution of diverse competitive strains of mesorhizobia suboptimal in symbiotic nitrogen fixation on the pasture legume Biserrula pelecinus L. Environmental Microbiology. 2007 Oct;9(10):2496–511.

40. Sullivan JT, Ronson CW. Evolution of rhizobia by acquisition of a 500-kb symbiosis island that integrates into a phe-tRNA gene. Proc Natl Acad Sci U S A. 1998 Apr 28;95(9):5145–9.

41. Kehlet-Delgado H, Montoya AP, Jensen KT, Wendlandt CE, Dexheimer C, Roberts M, et al. The evolutionary genomics of adaptation to stress in wild rhizobium bacteria. Proceedings of the National Academy of Sciences. 2024 Mar 26;121(13):e2311127121.

42. Porter SS, Rice KJ. Trade-Offs, Spatial Heterogeneity, and the Maintenance of Microbial Diversity. Evolution. 2013;67(2):599–608.

43. Porter SS, Chang PL, Conow CA, Dunham JP, Friesen ML. Association mapping reveals novel serpentine adaptation gene clusters in a population of symbiotic Mesorhizobium. ISME J. 2017 Jan;11(1):248–62.

44. Porter SS, Faber-Hammond J, Montoya AP, Friesen ML, Sackos C. Dynamic genomic architecture of mutualistic cooperation in a wild population of Mesorhizobium. The ISME Journal [Internet]. 2018 Sept 14; Available from: 10.1038/s41396-018-0266-y

45. Montoya AP, Kehlet-Delgado H, Jensen KT, Griffitts JS, Porter SS. Mobile genetic elements shape the evolution of wild bacterial pangenomes in the face of biotic and abiotic selective pressures in the soil [Internet]. bioRxiv; 2025 [cited 2025 Nov 13]. p. 2025.10.22.684009. Available from: https://www.biorxiv.org/content/10.1101/2025.10.22.684009v1

46. Somasegaran P, Hoben HJ. Handbook for Rhizobia: Methods in Legume-Rhizobium Technology [Internet]. New York: Springer-Verlag; 1994 [cited 2021 Apr 14]. Available from: https://www.springer.com/us/book/9781461383772

47. Wendlandt CE, Helliwell E, Roberts M, Nguyen KT, Friesen ML, von Wettberg E, et al. Decreased coevolutionary potential and increased symbiont fecundity during the biological invasion of a legume-rhizobium mutualism. Evolution. 2021;75(3):731–47.

48. Bañuelos-Vazquez LA, Castellani LG, Luchetti A, Romero D, Tejerizo GAT, Brom S. Role of plant compounds in the modulation of the conjugative transfer of pRet42a. PLOS ONE. 2020 Aug 26;15(8):e0238218.

49. Ling J, Wang H, Wu P, Li T, Tang Y, Naseer N, et al. Plant nodulation inducers enhance horizontal gene transfer of Azorhizobium caulinodans symbiosis island. Proc Natl Acad Sci U S A. 2016 Nov 29;113(48):13875–80.

50. Darling AE, Mau B, Perna NT. progressiveMauve: Multiple Genome Alignment with Gene Gain, Loss and Rearrangement. PLOS ONE. 2010 June 25;5(6):e11147.

51. Jain C, Rodriguez-R LM, Phillippy AM, Konstantinidis KT, Aluru S. High throughput ANI analysis of 90K prokaryotic genomes reveals clear species boundaries. Nat Commun. 2018 Nov 30;9(1):5114.

52. Grant JR, Enns E, Marinier E, Mandal A, Herman EK, Chen C yu, et al. Proksee: in-depth characterization and visualization of bacterial genomes. Nucleic Acids Research. 2023 July 5;51(W1):W484–92.

53. Montoya AP, Wendlandt CE, Benedict AB, Roberts M, Piovia-Scott J, Griffitts JS, et al. Hosts winnow symbionts with multiple layers of absolute and conditional discrimination mechanisms. Proceedings of the Royal Society B: Biological Sciences. 2023 Jan 4;290(1990):20222153.

54. Vincent JM. A Manual for the Practical Study of Root-nodule Bacteria. [Published for the] International Biological Programme [by] Blackwell Scientific; 1970. 202 p.

55. Regus JU, Quides KW, O’Neill MR, Suzuki R, Savory EA, Chang JH, et al. Cell autonomous sanctions in legumes target ineffective rhizobia in nodules with mixed infections. American Journal of Botany. 2017;104(9):1299–312.

56. Westhoek A, Field E, Rehling F, Mulley G, Webb I, Poole PS, et al. Policing the legume-Rhizobium symbiosis: a critical test of partner choice. Scientific Reports. 2017 May 3;7(1):1419.

57. Schneider CA, Rasband WS, Eliceiri KW. NIH Image to ImageJ: 25 years of image analysis. Nat Methods. 2012 July;9(7):671–5.

58. R Core Team. R: A language and environment for statistical computing. R Foundation for Statistical Computing, Vienna, Austria. 2022; Available from: https://www.R-project.org/.

59. Hartig F. DHARMa: Residual Diagnostics for Hierarchical (Multi-Level / Mixed) Regression Models [Internet]. 2016 [cited 2024 Oct 23]. p. 0.4.7. Available from: https://CRAN.Rproject.org/package=DHARMa

60. Bates D, Mächler M, Bolker B, Walker S. Fitting Linear Mixed-Effects Models Using lme4. Journal of Statistical Software. 2015 Oct 7;67:1–48.

61. Brooks ME, Kristensen K, Benthem KJ van, Magnusson A, Berg CW, Nielsen A, et al. glmmTMB Balances Speed and Flexibility Among Packages for Zero-inflated Generalized Linear Mixed Modeling. The R Journal. 2017;9(2):378–400.

62. Heath KD. Intergenomic epistasis and coevolutionary constraint in plants and rhizobia. Evolution. 2010 May;64(5):1446–58.

63. Shaw RG, Shaw FH. Quantitative genetic study of the adaptive process. Heredity. 2014 Jan;112(1):13–20.

64. Canty A, Ripley B. boot: Bootstrap R (S-Plus) Functions. R package version. 2016 Jan 1;1:3–18.

65. Lenth RV. emmeans: Estimated Marginal Means, aka Least-Squares Means [Internet]. 2017 [cited 2024 Oct 23]. p. 1.10.5. Available from: https://CRAN.Rproject.org/package=emmeans

66. Pasch B, Bolker BM, Phelps SM. Interspecific Dominance Via Vocal Interactions Mediates Altitudinal Zonation in Neotropical Singing Mice. The American Naturalist. 2013 Nov;182(5):E161–73.

67. Batstone RT. Genomes within genomes: nested symbiosis and its implications for plant evolution. New Phytologist [Internet]. 2022 [cited 2025 Apr 3];234(1):28–34. Available from: https://onlinelibrary.wiley.com/doi/abs/10.1111/nph.17847

68. Haudiquet M, de Sousa JM, Touchon M, Rocha EPC. Selfish, promiscuous and sometimes useful: how mobile genetic elements drive horizontal gene transfer in microbial populations. Philosophical Transactions of the Royal Society B: Biological Sciences. 2022 Aug 22;377(1861):20210234.

69. Geddes BA, Kearsley J, Morton R, diCenzo GC, Finan TM. Chapter Eight - The genomes of rhizobia. In: Frendo P, Frugier F, Masson-Boivin C, editors. Advances in Botanical Research [Internet]. Academic Press; 2020 [cited 2024 Oct 20]. p. 213–49. (Regulation of Nitrogen-Fixing Symbioses in Legumes; vol. 94). Available from: https://www.sciencedirect.com/science/article/pii/S0065229619300916

70. Westhoek A, Clark LJ, Culbert M, Dalchau N, Griffiths M, Jorrin B, et al. Conditional sanctioning in a legume– *Rhizobium* mutualism. Proc Natl Acad Sci USA. 2021 May 11;118(19):e2025760118.

71. Hardiman CA, Weingarten RA, Conlan S, Khil P, Dekker JP, Mathers AJ, et al. Horizontal Transfer of Carbapenemase-Encoding Plasmids and Comparison with Hospital Epidemiology Data. Antimicrobial Agents and Chemotherapy. 2016 July 22;60(8):4910–9.

72. Allard N, Collette A, Paquette J, Rodrigue S, Côté JP. Systematic investigation of recipient cell genetic requirements reveals important surface receptors for conjugative transfer of IncI2 plasmids. Commun Biol. 2023 Nov 16;6(1):1–11.

73. Pérez-Mendoza D, de la Cruz F. Escherichia coli genes affecting recipient ability in plasmid conjugation: Are there any? BMC Genomics. 2009 Feb 9;10(1):71.

74. Remigi P, Zhu J, Young JPW, Masson-Boivin C. Symbiosis within Symbiosis: Evolving Nitrogen-Fixing Legume Symbionts. Trends in Microbiology. 2016 Jan;24(1):63–75.

75. Wardell GE, Hynes MF, Young PJ, Harrison E. Why are rhizobial symbiosis genes mobile? Philosophical Transactions of the Royal Society B: Biological Sciences. 2022 Jan 17;377(1842):20200471.

76. Colombi E, Perry BJ, Sullivan JT, Bekuma AA, Terpolilli JJ, Ronson CW, et al. Comparative analysis of integrative and conjugative mobile genetic elements in the genus Mesorhizobium. Microbial Genomics [Internet]. 2021 Oct 4 [cited 2022 Aug 30];7(10). Available from: https://www.microbiologyresearch.org/content/journal/mgen/10.1099/mgen.0.000657

77. Ramsay JP, Sullivan JT, Stuart GS, Lamont IL, Ronson CW. Excision and transfer of the Mesorhizobium loti R7A symbiosis island requires an integrase IntS, a novel recombination directionality factor RdfS, and a putative relaxase RlxS. Molecular Microbiology. 2006;62(3):723–34.

78. Ramsay JP, Sullivan JT, Jambari N, Ortori CA, Heeb S, Williams P, et al. A LuxRI-family regulatory system controls excision and transfer of the Mesorhizobium loti strain R7A symbiosis island by activating expression of two conserved hypothetical genes. Molecular Microbiology. 2009;73(6):1141–55.

79. Ramsay JP, Major AS, Komarovsky VM, Sullivan JT, Dy RL, Hynes MF, et al. A widely conserved molecular switch controls quorum sensing and symbiosis island transfer in Mesorhizobium loti through expression of a novel antiactivator. Molecular Microbiology. 2013;87(1):1–13.

80. diCenzo GC, MacLean AM, Milunovic B, Golding GB, Finan TM. Examination of Prokaryotic Multipartite Genome Evolution through Experimental Genome Reduction. PLOS Genetics. 2014 Oct 23;10(10):e1004742.

81. Perry BJ, Sullivan JT, Colombi E, Murphy RJT, Ramsay JP, Ronson CW. Symbiosis islands of Loteae-nodulating Mesorhizobium comprise three radiating lineages with concordant nod gene complements and nodulation host-range groupings. Microb Genom. 2020 Aug 26;6(9):mgen000426.

82. Johnson CN, Sheriff EK, Duerkop BA, Chatterjee A. Let Me Upgrade You: Impact of Mobile Genetic Elements on Enterococcal Adaptation and Evolution. Journal of Bacteriology. 2021 Oct 12;203(21):10.1128/jb.00177-21.

83. Rocha EPC, Bikard D. Microbial defenses against mobile genetic elements and viruses: Who defends whom from what? PLOS Biology. 2022 Jan 13;20(1):e3001514.

84. Hill YJ, Kohlmeier MG, Agha Amiri A, O’Hara GW, Terpolilli JJ. Evolution of novel Mesorhizobium genospecies that competitively and effectively nodulate Cicer arietinum following inoculation with the Australian commercial inoculant strain M. ciceri CC1192. Plant Soil [Internet]. 2025 Feb 1 [cited 2025 Apr 12];507(1):397–415. Available from: 10.1007/s11104-024-06739-y

85. Riaz MR, Sosa Marquez I, Lindgren H, Levin G, Doyle R, Romero MC, et al. Mobile gene clusters and coexpressed plant–rhizobium pathways drive partner quality variation in symbiosis. Proceedings of the National Academy of Sciences [Internet]. 2025 Aug 5 [cited 2025 July 31];122(31):e2411831122. Available from: https://www.pnas.org/doi/10.1073/pnas.2411831122

86. Hirsch AM, Wilson KJ, Jones JD, Bang M, Walker VV, Ausubel FM. Rhizobium meliloti nodulation genes allow Agrobacterium tumefaciens and Escherichia coli to form pseudonodules on alfalfa. Journal of Bacteriology. 1984 June;158(3):1133–43.

87. Martínez E, Palacios R, Sánchez F. Nitrogen-fixing nodules induced by Agrobacterium tumefaciens harboring Rhizobium phaseoli plasmids. Journal of Bacteriology. 1987 June;169(6):2828–34.

88. Rogel MA, Hernández-Lucas I, Kuykendall LD, Balkwill DL, Martinez-Romero E. Nitrogen-Fixing Nodules with Ensifer adhaerens Harboring Rhizobium tropici Symbiotic Plasmids. Applied and Environmental Microbiology. 2001 July;67(7):3264–8.

89. Bean EL, Herman C, Anderson ME, Grossman AD. Biology and engineering of integrative and conjugative elements: Construction and analyses of hybrid ICEs reveal element functions that affect species-specific efficiencies. PLOS Genetics. 2022 May 18;18(5):e1009998.

90. Asnicar F, Thomas AM, Beghini F, Mengoni C, Manara S, Manghi P, et al. Precise phylogenetic analysis of microbial isolates and genomes from metagenomes using PhyloPhlAn 3.0. Nat Commun. 2020 May 19;11(1):2500.

91. Stamatakis A. RAxML version 8: a tool for phylogenetic analysis and post-analysis of large phylogenies. Bioinformatics. 2014 May 1;30(9):1312–3.

