## Supplemental Information for "The evolutionary genomics of incipient endosymbiosis in wild rhizobia bacteria"

This file includes supplemental methods and results, including Figures S1, S2, and S3, and Tables S2, S5, S6, S7 and S8. Additional supplementary tables (Tables S1, S3, and S4) can be found in the supplemental excel file.

### I. Plasmid construction to track the *Mesorhizobium* symbiosis island (SI)

A plasmid was designed to track the symbiosis island during conjugative transfer and contains selectable markers and a region of homology with the symbiosis island (SI) to enable the plasmid to integrate site specifically into the SI via homologous recombination. We targeted a 500 bp region at the end of the *nifA* gene (including the stop codon and intergenic region after the gene) in the SI for the region of homology since it is a highly conserved operon. However, multiple sequence alignments of potential *Mesorhizobium* SI donors showed there were still several single nucleotide polymorphisms in the targeted region. Therefore, two SI tracking plasmids (pKJ116, pKJ117) were designed from the same plasmid (pJG1108) but have slightly different variations of the *nifA* gene region of homology.

The SI tracking plasmid was first introduced via bacterial transformation into specialized *E. coli* strain MFDpir (1) which is auxotrophic for DAP (diaminopimelic acid- a lysine precursor). MFDpir isolates with either reporter plasmid (pKJ116 or pKJ117) (Figure S1) were grown at 37°C overnight on (LB) agar (2) supplemented with 50 µg/mL DAP and 30 µg/mL Kanamycin. *Mesorhizobium* were grown at 28°C for 5 days on Tryptone Yeast (TY) agar (3). Approximately equal amounts of MFDpir and *Mesorhizobium* cells were re-suspended in LB broth or TY broth, respectively, and 12 µL of each donor and recipient were mixed in separate 1.5 mL microcentrifuge tubes. 20µL of each bi-parental mating mixture was spot plated on TY agar with 50 µg/mL DAP and incubated at 28°C for 6-24 hours prior to selecting for *Mesorhizobium* SI donor transconjugants on TY agar with 100 µg/mL neomycin (Nm). Then *Mesorhizobium* SI donor transconjugants were cryopreserved after testing on X-Gluc media, where SI donors containing the SI tracking plasmid turn blue due to the presence of the *gusA* gene. To generate selectable recipient strains, *Mesorhizobium* strains that lack the SI were selected for spontaneous dual antibiotic resistance by culturing initially sensitive isolates first on streptomycin (Sm; 200 µg/mL) until a resistant mutant arose, then on rifampicin (Rf; 100 µg/mL) until a resistant mutant arose. Thus, we generated donor strains that are Nm resistant and blue in the presence of X-Gluc, as well as recipient cells that are Sm and Rf resistant and colorless in the presence of X-Gluc. Novel transconjugants that result from the transmission of the SI from donor to recipient are Nm, Sm, and Rf resistant and blue in the presence of X-Gluc.

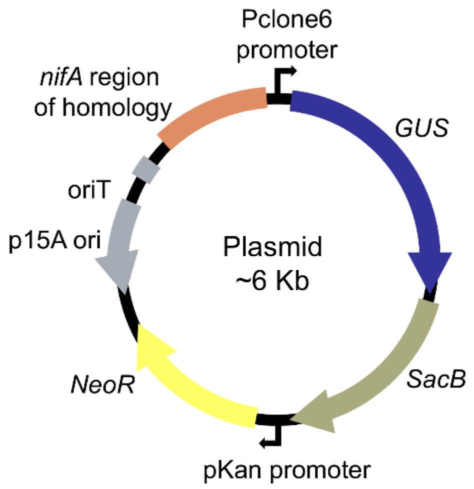


**Figure S1. Genetic map of the SI tracking plasmid that inserts into the symbiosis island of wild *Mesorhizobium* strains.** The *Mesorhizobium* SI is marked with a plasmid containing selectable marker genes Neomycin (*NeoR*) resistance, *gusA* (GUS), *SacB* (not used in this experiment), and a region of homology with the *nifA* gene to enable the plasmid to integrate site specifically into the intergenic region after the *nifA* gene via homologous recombination. Two SI tracking plasmids, pKJ116 (6039 bp) and pKJ117 (6034 bp) have different variations of the *nifA* region of homology. The map also indicates locations of the promoters, *Pclone6* and *pKan*, the cis-acting mobilization determinant (*oriT*), and the *E. coli* replication origin, *p15A ori*.

### II. SI transmission

Mating assays had 15 donors and 6 recipients and each strain alone as negative controls. In assay 2 we included a positive control of a donor-recipient combination used in assay 1 that also had successful SI transfer in pilot experiments. To prepare donor and recipient cells for the mating assay, we estimated cell density as $OD_{600} \times 5.8 \times 10^7$ = CFU $mL^{-1}$ (4) and resuspended cells to 6 x $10^8$ CFU $mL^{-1}$. Mating spots consisting of 10 µL of the donor culture, 10 µL of the recipient culture, and 10 µL of plant root extracts were plated on TY agar and incubated for 40 hours (4 spots per 60 mm plate). Negative controls consisting of 10 µL of each strain and 10 µL of plant root extracts were spot plated along with the matings. At the end of each mating assay, the entire spot was gathered with a sterile pestle and suspended in 350 µL of TY broth with Nm. Then each culture (mating spots, and negative and positive controls) was plated on selection plates, each with 3 replicates. There was a total of 804 selection plates (672 round 100 mm plates and 132 square 120 mm plates). Across all successful matings, there was an average of $1.7 \times 10^4$ (95% CI: [$1.1 \times 10^4$, $2.2 \times 10^4$], SE = $2.6 \times 10^3$) transconjugants per mating spot.

#### *Confirmatory PCR*

A subset of the transconjugants were tested using colony PCR to confirm SI presence and the identity of the recipient genotype. To confirm the identity of SI gain transconjugants, we used draft genome data from Kehlet-Delgado et al. (2024) and Porter et al. (2017) to design genotype-specific primers to amplify a section of one gene unique to each recipient strain (5–7). Following Ferrandis-Vila et al. (2022), we defined the pangenome of all strains in this experiment using Roary (v 3.12.0) (8) and used the gene presence absence table to identify a unique gene in each recipient. Primers were designed using Primer3Plus (v 3.3.0) (9). However, the primers designed for recipient R06 were not successful in PCR assays but the primers and PCR assays for the 5 other recipients worked (Table S2). In addition, we used established primer sets in PCR assays for donors and SI gain transconjugants to amplify part of the *nodA* gene in the SI to confirm SI presence (10) (Table S2). In colony PCR, cells from single colonies were suspended in 10 μL nuclease-free water and 1 μL served as template DNA in the 10 μL PCR. This cycled in a thermocycler with parameters set to 3 min at 95°C, followed by 35 cycles of 20 s at 92°C for denaturation, 20 s at 56°C (*nodA* primers) or 68°C (genotype-specific primers) for annealing, and 2 min at 68°C for extension, then an additional 3 min at 68 °C. PCR products were visualized under UV light, on a 1.2% agarose gel pre-stained with SafeView Classic.

**Table S2. Primers (Oligonucleotides) used in this study.** The *nodA* primer set was designed previously (10). The genotype specific primers were designed for this study and are named as the “wildtype recipient strain name_locus tag of unique gene in the original draft genome” (6,7).

| Primers | Sequence | Description |
|---|---|---|
| nodA_69F | CGCCGAGTTCTTTCGTGATA | Forward primer to amplify part of nodA gene. Use with nodA_390R |
| nodA_390R | TCCGAACCTCTCAACATGATTC | Reverse primer to amplify part of nodA gene. Use with nodA_69F |
| C380A_02022F | AAAGCCCGAGGTAGTTTGGT | Forward primer to amplify part of unique gene in recipient R01. Use with C380A_02022R |
| C380A_02022R | AAGCCCAAGCAGAGAAACAA | Reverse primer to amplify part of unique gene in recipient R01. Use with C380A_02022F |
| C398B_04160F | CTAAAGCCAAGCACCCTCTG | Forward primer to amplify part of unique gene in recipient R02. Use with C398B_04160R |
| C398B_04160R | TTGATCTGCTTGACGACGAC | Reverse primer to amplify part of unique gene in recipient R02. Use with C398B_04160F |
| C403B_00763F | CCAGAGTTCTGGCTTTACCG | Forward primer to amplify part of unique gene in recipient R03. Use with C403B_00763R |
| C403B_00763R | CTGGCGTATATGGGCGTACT | Reverse primer to amplify part of unique gene in recipient R03. Use with C403B_00763F |
| C432A_00044F | CTGGAGGGCTACAGAGCATC | Forward primer to amplify part of unique gene in recipient R04. Use with C432A_00044R |
| C432A_00044R | TCCTTCGTTTTGGAGATTGG | Reverse primer to amplify part of unique gene in recipient R04. Use with C432A_00044F |
| C565B_01124F | CCAGATACAGGGCGATCCTA | Forward primer to amplify part of unique gene in recipient R05. Use with C565B_01124R |
| C565B_01124R | AACAACTGCCCCTGAATGTC | Reverse primer to amplify part of unique gene in recipient R05. Use with C565B_01124F |

**Table S5. Donor genotype, recipient genotype and their interaction impact symbiosis island transfer**. Percent of the variance for each component and likelihood ratio test $\chi^2$ statistics for random effects in the number of transconjugants per mating variance component model. (*** P < 0.001, ** P < 0.01, * P < 0.05). 95% confidence interval (CI) from 10,000 nonparametric bootstrap replicates of the model is also shown.

| Number of transconjugants per mating | | | |
|---|---|---|---|
| **Component** | **% Variance** | $\chi^2$ | **CI** |
| Donor | 48.48 | 94.03*** | [45.37 - 51.86] |
| Recipient | 24.98 | 78.67*** | [22.88 - 27.59] |
| Donor x Recipient | 26.52 | 632.97*** | [22.53 - 29.28] |

### III. Symbiotic function of SI gain transconjugants

We extracted and cultured 92 nodules from five blocks to determine the number of colony forming units (CFUs) per nodule (*CFU/nodule*). However, 25% of nodules failed to yield viable rhizobium colonies. We also excluded four nodules that did not culture well because the size of these nodules were as large or larger than other replicate nodules but the CFU data was several orders of magnitude lower. This poor nodule culturing may suggest that nodules were beginning to senesce (plants were ~15 weeks old and some were beginning to flower). In total, 65 nodules contributed data to *CFU/nodule*, with less than 3 biological replicates for 2/5 donors and 4/15 SI gain transconjugants. Therefore, we only use the *CFU/nodule* data for the transconjugant variance component model (48 nodules) (Figure S2). Donors formed greater CFU per nodule on average ($6.1 \times 10^6$; 95% CI: [$1.2 \times 10^6$, $3.2 \times 10^7$], SE = $4.8 \times 10^6$) compared to the transconjugants ($2.6 \times 10^5$; 95% CI: [$9.4 \times 10^4$, $7.1 \times 10^5$], SE = $1.2 \times 10^5$).

We also used a linear model with log transformed CFU per nodule predicted by nodule width. Here, we found nodule width is positively correlated with *CFU/nodule* ($r$ = 0.33, $P$ = 0.007; Figure S3) and the amount of variance explained by genetic components do not differ among nodule width and *CFU/nodule* (Table S6; Figure 4e.f; Figure S2). Therefore, we expect the other *nodule width* model results would be similar for the number of progeny within nodules.

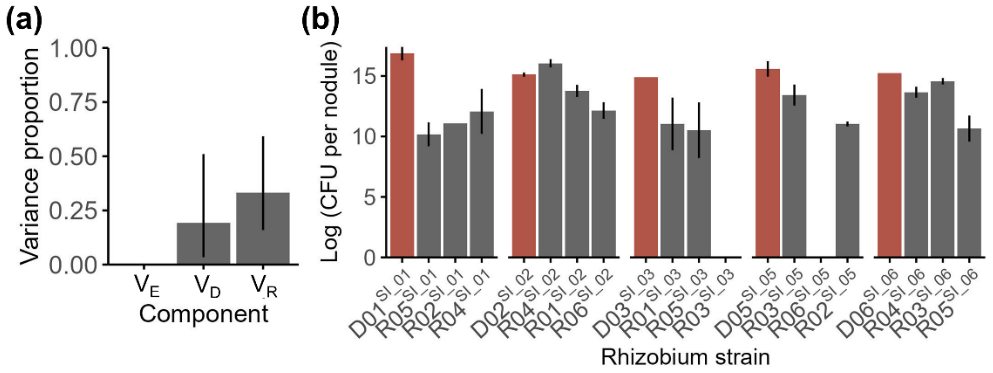


**Figure S2: The proportion of variance explained by genetic components does not differ between the number of rhizobium progeny per nodule and nodule width.** The percent of phenotypic variance in (a) the number of colony forming units (CFUs) per nodule. Bars show variance estimates for experimental block ($V_E$), donor-SI genotype ($V_D$) and recipient genotype ($V_R$) and 95% confidence intervals. No confidence interval is shown for block because it did not contain the original estimate of zero variance. (b) CFU per nodule for SI donors and SI gain transconjugants. Within each donor SI type (SI_01, SI_02, SI_03, SI_05, and SI_06), the donor is shown first in red (D01, D02, D03, D05 and D06), followed by the three transconjugants (named by recipient genotype R01 - R06). Bars are genotypic means +/- standard error.

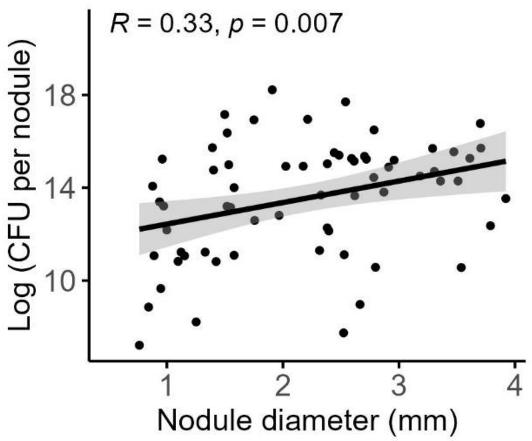


**Figure S3. Nodule width is positively correlated with the number of progeny per nodule.** Filled circles show the mean values of CFU and nodule width for each cultured nodule (n= 65). Values shown in the plot are the Pearson correlation coefficient and p-value. Black line, linear regression. Grey shading, standard error.

**Table S6. Symbiosis island function following transfer is impacted by the SI donor and recipient genotypes**. Percent of the variance and likelihood ratio test $\chi^2$ statistics for random effects in host fitness (shoot mass, percent of nitrogen in leaf tissue) and symbiont fitness (nodule width, number of nodules, number of N-fixing nodules, and CFU per nodule) variance component models. (*** P < 0.001, ** P < 0.01, * P < 0.05).

| Component | Donor | | Recipient | | Block | | Residual |
|---|---|---|---|---|---|---|---|
| | % var | χ2 | % var | χ2 | % var | χ2 | % var |
| **Phenotype** | | | | | | | |
| Shoot mass | 6.9 | 16.9*** | 4.1 | 8.5** | 60.8 | 138.1*** | 28.2 |
| Leaf %N | 55.5 | 74.7*** | 43.9 | 27.9*** | 0.66 | 0.08 | 39.9 |
| Nodule number | 43.8 | 25.9*** | 46.4 | 15.6*** | 9.9 | 3.5 | |
| N-fixing nodule number | 23.1 | 127.4*** | 76.2 | 108.4*** | 0.7 | 5.2* | |
| Nodule width | 38.5 | 81.12*** | 31.1 | 53.92*** | 3.9 | 8.15*** | 26.5 |
| CFU/nodule | 33.2 | 5.87* | 19.3 | 8.12** | 0 | 0 | 47.5 |

**Table S7. Statistical tests of the effect of eco-eco factors on SI function for host plant and rhizobium symbiotic fitness.** We used GLMM models with phylogenetic and geographic distance, soil type and host plant species as covariates and with donor genotype, recipient genotype, their interaction, and block as random effects. However, random effects were removed if the model was overparameterized: donor-SI was removed from the leaf tissue percent nitrogen model, and the interaction of donor-SI and recipient genotype was removed from the N-fixing nodule number model. The table displays likelihood ratio test χ2 values for each model, with significance of each measure indicated (*** P < 0.001, ** P < 0.01, * P < 0.05).

| Trait | Shoot mass | Leaf %N | Nodule width | N-fixing nodule number | Nodule number |
|---|---|---|---|---|---|
| **Predictor** | | | | | |
| Phylogenetic distance | 7.91** | 14.6*** | 16.19*** | 19.93*** | 0.664 |
| Soil ecotype | 5.56* | 6.45* | 5.11* | 19.30*** | 0.004 |
| Host species type | 0.37 | 0.94 | 0.69 | 0.0001 | 0.005 |
| Geographic distance | 0.85 | 0.79 | 0.03 | 1.53 | 0.112 |

**Table S8. Statistical tests of the effect of eco-eco factors on SI transmission.** The number of transconjugants per mating was used as the response variable in a GLMM with phylogenetic and geographic distance, host plant species and soil type as fixed effects and donor-SI genotype, recipient genotype, and their interaction as random effects. The table displays likelihood ratio test $\chi 2$ values, with significance of each measure indicated (*** $P < 0.001$, ** $P < 0.01$, * $P < 0.05$).

| Number of transconjugants per mating | |
|---|---|
| **Predictor** | $\chi^2$ |
| Phylogenetic distance | 0.56 |
| Geographic distance | 0.04 |
| Soil ecotype | 0.98 |
| Host species | 0.04 |

**Supplemental References**


1. Ferrières L, Hémery G, Nham T, Guérout AM, Mazel D, Beloin C, et al. Silent Mischief: Bacteriophage Mu Insertions Contaminate Products of Escherichia coli Random Mutagenesis Performed Using Suicidal Transposon Delivery Plasmids Mobilized by Broad-Host-Range RP4 Conjugative Machinery. Journal of Bacteriology. 2010 Dec 15;192(24):6418–27.

2. Bertani G. STUDIES ON LYSOGENESIS I.: The Mode of Phage Liberation by Lysogenic Escherichia coli. Journal of Bacteriology. 1951 Sept 1;62(3):293–300.

3. Somasegaran P, Hoben HJ. Handbook for Rhizobia: Methods in Legume-Rhizobium Technology [Internet]. New York: Springer-Verlag; 1994 [cited 2021 Apr 14]. Available from: https://www.springer.com/us/book/9781461383772

4. Wendlandt CE, Helliwell E, Roberts M, Nguyen KT, Friesen ML, von Wettberg E, et al. Decreased coevolutionary potential and increased symbiont fecundity during the biological invasion of a legume-rhizobium mutualism. Evolution. 2021;75(3):731–47.

5. Ferrandis-Vila M, Tiwari SK, Mamerow S, Semmler T, Ferrandis-Vila M, Tiwari SK, et al. Using unique ORFan genes as strain-specific identifiers for Escherichia coli. BMC Microbiology. 2022 May 18;22(1):135.

6. Kehlet-Delgado H, Montoya AP, Jensen KT, Wendlandt CE, Dexheimer C, Roberts M, et al. The evolutionary genomics of adaptation to stress in wild rhizobium bacteria. Proceedings of the National Academy of Sciences. 2024 Mar 26;121(13):e2311127121.

7. Porter SS, Chang PL, Conow CA, Dunham JP, Friesen ML. Association mapping reveals novel serpentine adaptation gene clusters in a population of symbiotic Mesorhizobium. ISME J. 2017 Jan;11(1):248–62.

8. Page AJ, Cummins CA, Hunt M, Wong VK, Reuter S, Holden MTG, et al. Roary: rapid large-scale prokaryote pan genome analysis. Bioinformatics. 2015 Nov 15;31(22):3691–3.

9. Untergasser A, Cutcutache I, Koressaar T, Ye J, Faircloth BC, Remm M, et al. Primer3—new capabilities and interfaces. Nucleic Acids Research. 2012 June 21;40(15):e115.

10. Porter SS, Faber-Hammond J, Montoya AP, Friesen ML, Sackos C. Dynamic genomic architecture of mutualistic cooperation in a wild population of Mesorhizobium. The ISME Journal [Internet]. 2018 Sept 14; Available from: https://doi.org/10.1038/s41396-018-0266-y
